# Genomic variation associated with endosymbiont shuffling and areal growth in the endangered elkhorn coral, *Acropora palmata*

**DOI:** 10.64898/2026.08.14.744775

**Authors:** Ruiqi Li, Holland Elder, Grace McDermott, Sibelle O’Donnell, Courtney Klepac, Maria Ruggeri, Sophia Lee, Wyatt C. Million, Zachary Craig, Dakotah Merck, Erinn M. Muller, Carly D. Kenkel

## Abstract

Biodiversity losses continue to outpace traditional management, underscoring the need to understand adaptive capacity and the potential for interventions to increase fitness under climate change. We undertook a genome-wide association study on 156 *Acropora palmata* genets to investigate the genomic basis of areal growth, endosymbiont association, and thermal tolerance. Seven peaks on chromosomes 1, 3 and 14 were associated with endosymbiont shuffling and two peaks on chromosome 4 were associated with areal growth. As variants were located in non-coding regions we incorporated additional data from an independent field-transplant experiment to investigate their relationship with patterns of gene expression. Intersection of these datasets implicated melanocortin-like receptor activity and Ran GTPase activating protein 1 in endosymbiont composition and surface area growth, respectively. Results indicate that growth and endosymbiont associations may represent more viable intervention targets than temperature tolerance and highlight the need to better understand the role of non-coding variation in basic biology and development of restoration interventions.

## INTRODUCTION

The ability to predict phenotype from genomic data has been recognized as the next frontier in conservation management, informing development of interventions including live- and cryo-banking, selective breeding, and assisted gene flow^1,2^. Such intervention strategies are increasingly relevant in this era of rapid global change as biodiversity losses continue to outpace traditional management approaches^3,4^. Identifying genomic markers of key phenotypes in reef-building coral is of particular urgency^5,6^ given their global ecological significance as foundational ecosystem builders and high risk of extinction due to global climate change^7,8^. Studies of temperature tolerance traits have dominated the field to date, with the emerging consensus that coral thermal tolerance is polygenic, resulting from allelic variation at many small-effect loci^5,9–11^. This means that while many adaptive solutions are possible, the large number of loci, variation in their relative contribution to phenotype and potential interactions, as well as fluctuating contributions of individual alleles over space and time (e.g. generations^12^) limit their predictive utility. Other key phenotypes, such as growth, pathogen resistance and symbiont association, may represent more viable intervention targets. While Vollmer *et al*. (2023)^6^ recently identified just 10 genomic regions in *Acropora cervicornis* associated with resistance to white band disease in support of this promise, the genomic basis of key performance traits apart from thermal tolerance remains underexplored.

Identification of host loci associated with algal endosymbiont community composition represents a particularly critical knowledge gap given the well-established role of symbionts in coral thermal tolerance. Genome-wide association studies (GWAS) have found that a significant portion of the variation in coral thermal tolerance is actually explained by the dominant symbiotic partner^5,9,11^. For example, in a genomic prediction model for *Acropora millepora*, Fuller et al. (2020) found that almost 20% of the variation in bleaching was attributable to the proportion *Durusdinium* spp. hosted by a coral genet, whereas the cumulative effect of host polygenic score yielded only an additional 2.6% in predictive power. If the symbiotic associations themselves can be more reliably predicted by host genotype, rather than the phenotypes they contribute to, this may represent a more viable target for selection. Encouragingly, symbiont specificity appears heritable^13^ and Hawaiian *Montipora capitata*, for example, exhibit a stable polymorphism for symbiont preference, with individual coral remaining largely *Cladocopium* or *Durusdinium* dominated even through bleaching events^14–16^. Yet coral endosymbiont associations are complex and also environment-dependent. The majority of coral species must acquire their symbionts from the environment early in development^17^. This initial colonization is generally promiscuous followed by subsequent winnowing to a (mostly) species-specific association in adulthood^18^. Subsequent changes in community composition via shuffling^19^(i.e. changes in relative proportions of existing symbionts) or switching^20^ (i.e. acquisition of novel symbionts) can also occur in response to environmental variation. *Durusdinium* spp. endosymbionts are known to confer some of the highest thermal tolerances^21^, but metabolic trade-offs, such as reduced carbon translocation to their coral hosts, are thought to limit their universal acquisition^22,23^, especially outside periods of thermal stress. While some progress has been made in identifying genes associated with symbiotic state (i.e. symbiotic vs. aposymbiotic coral^24^), the genomic basis of partner preference (if any) remains unresolved.

While thermal tolerance is important for surviving both seasonal temperature fluctuations and extreme anomalies, coral growth is equally important for the functional significance of reefs. Growth of coral builds the three-dimensional structure of reefs which generates habitat for other organisms^25^, directly supporting fisheries and reef tourism^26^ and protecting local coastlines^27^. Local environmental variation can either promote or constrain patterns and rates of growth, but growth also varies among and within species^28–31^ indicating a host genetic component. In support of this latter observation, studies across species have confirmed growth is a heritable trait^32^, although the contributions of the endosymbiont remain unresolved^33^. In parallel, functional genomic and proteomic studies have begun to elucidate the genes involved in coral skeletal growth (∼calcification or biomineralization), including identifying novel cassettes of genes such as the coral acid-rich proteins (CARPs) which appear to be unique to calcifying coral^34–36^. Yet, specific genomic variants associated with individual variation in growth have yet to be identified.

We undertook a genome-wide association study to investigate the genomic basis of areal growth, symbiont association and thermal tolerance traits using 156 genets of the endangered elkhorn coral, *Acropora palmata*, of which 152 were sexually produced offspring of crosses among wild founder coral that have been used in reef restoration operations in Florida since 2018^37^ (Fig. 1). Genets were raised in captivity in a common garden nursery, thus limiting the effect of environmental history on variation in phenotype and enabling independent replication of genets across treatments using asexually produced ramets (∼independent fragments of the same genotype). In addition to collecting data on growth and thermal tolerance in response to a one-month long temperature stress exposure (Fig. 1B,C), we also took advantage of a serendipitous symbiont shuffling event to quantify changes in dominant endosymbiont associations among genets. As variants associated with growth and retention of dominant endosymbiont type were not located in coding regions we hypothesized that a change in gene regulation, rather than gene function^38^, could be involved. We therefore incorporated additional growth and gene expression data from an independent field-transplant of a subset of five genets to nine reef sites in the lower Florida Keys in which shuffling of endosymbiont communities was also observed^39^ (Fig. 1D-G). Genomic regions highlighted by both analyses were considered to be strong candidates underpinning trait variation (Fig. 1G).

**Figure 1.**
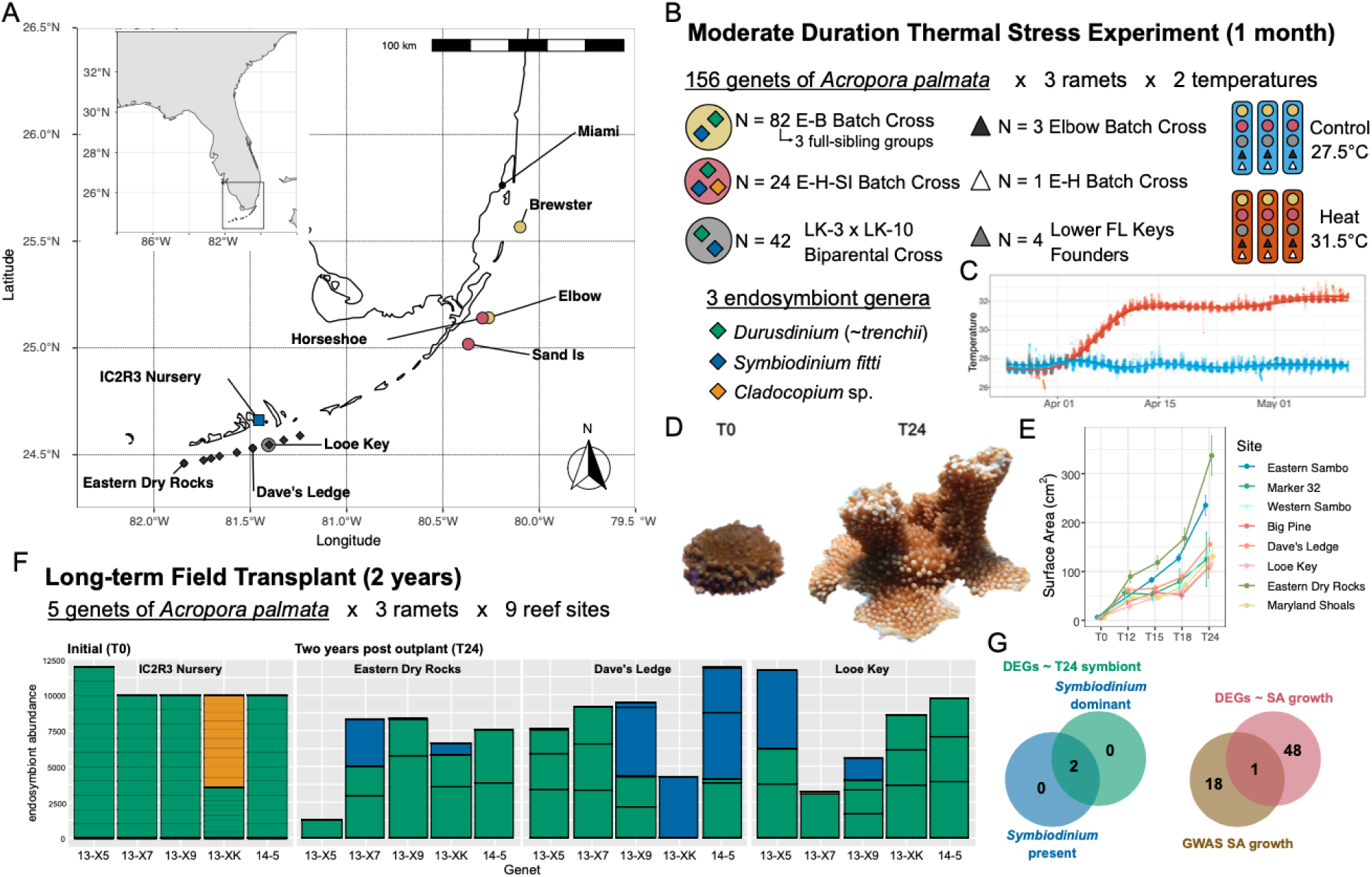
Experimental thermal stress, GWAS, and field transplant design. (A) Map of parental source (colored circles) and transplant (diamonds) reef sites, including the location of the IC2R3 *ex situ* coral nursery. (B) Six ramets of each of 156 genets of *Acropora palmata,* hosting different ratios of dinoflagellate endosymbiont genera and derived from different batch crosses and field collections^37^, were subjected to a one-month long thermal stress experiment (N=3 ramets/treatment). (C) Temperature ramp profiles for replicate experimental raceways at IC2R3. (D) Example image of *A. palmata* from the IC2R3 nursery and 24 months post-transplant (T24). (E) Average surface area (+/- SEM) of outplants over time grouped by transplant destination. (F) In 2018, three replicate ramets of each of five *A. palmata* genets were transplanted to nine different reef sites in the Lower Florida Keys for two years. Both algal endosymbiont community composition^39^ and global gene expression patterns (G) were profiled before and after transplantation and related to changes in symbiont community composition and surface area growth. Venn diagrams indicate the number of significantly differentially expressed genes satisfying each selection criterion.

## RESULTS

### Genet-specific differences in algal endosymbiont associations persist under thermal stress

Captive-bred *A. palmata* reared in Mote Marine Laboratory’s land-based facility (IC2R3) are known to associate with multiple algal partners and were historically dominated by the heterologous symbiont, *Durusdinium* (∼*trenchii*)^39,40^. Relative quantification of symbiont genera using qPCR revealed that although most ramets of our 156 experimental genets remained dominated by *Durusdinium trenchii*, some had begun to transition to *Symbiodinium* (∼*fitti*), while a few genets exhibited *Cladocopium* spp. dominance in line with prior observations^39,40^.

As algal endosymbiont composition can influence holobiont thermal tolerance^5,21^ as well as be influenced by thermal history^19,41^, and there is some evidence for a heritable host genetic influence on endosymbiont associations^13^, we tested for effects of genet, family origin (five full-sibling families derived from batch crosses^37^) and temperature treatment on dominant symbiont types using a series of Beta regression models. No significant interaction terms were detected for any model. Models incorporating fixed effects of genet nested within family and temperature treatment had the highest explanatory power overall (%*Durusdinium* ΔAIC=511.2, pseudo R^2^=0.88; %*Symbiodinium* ΔAIC=473.4, pseudo R^2^=0.87), indicating that symbiont associations were largely consistent across ramets of a genet (Fig. 2A,S1). Pairwise comparisons against a *Durusdinium* dominated reference genet (E-H-SI Genet 2, Fig. 2A) reinforced family-specific differences, where Biparental cross genets from Looe Key were almost exclusively *Durusdinium* dominated, with only one genet (AP331) out of 42 having significantly reduced abundances of *Durusdinium* (p=0.03) and correspondingly elevated abundances of *Symbiodinium* (p=0.055, Fig. 2A). Whereas 12 (out of 45 total, ∼27%) E-B batch cross full-sibling family A genets, 7 (out of 17 total, ∼41%) E-B batch full-sib family B genets, 6 (out of 18 total, ∼33%) E-B batch full-sib family C genets, and 11 (out of 24 total, ∼46%) E-H-SI batch cross genets had significantly reduced abundances of *Durusdinium* (p<0.05) relative to the reference genet (Fig. 2A). Temperature treatment also had a small but noticeable effect, with ramets in the heat treatment more likely to retain higher relative abundances of *Durusdinum*, by 2.5% on average, than those in control conditions (p<0.0001) and correspondingly hosting lower relative abundances of *Symbiodinium*, by 1.8% on average (p<0.0001, Fig. 2A).

**Figure 2.**
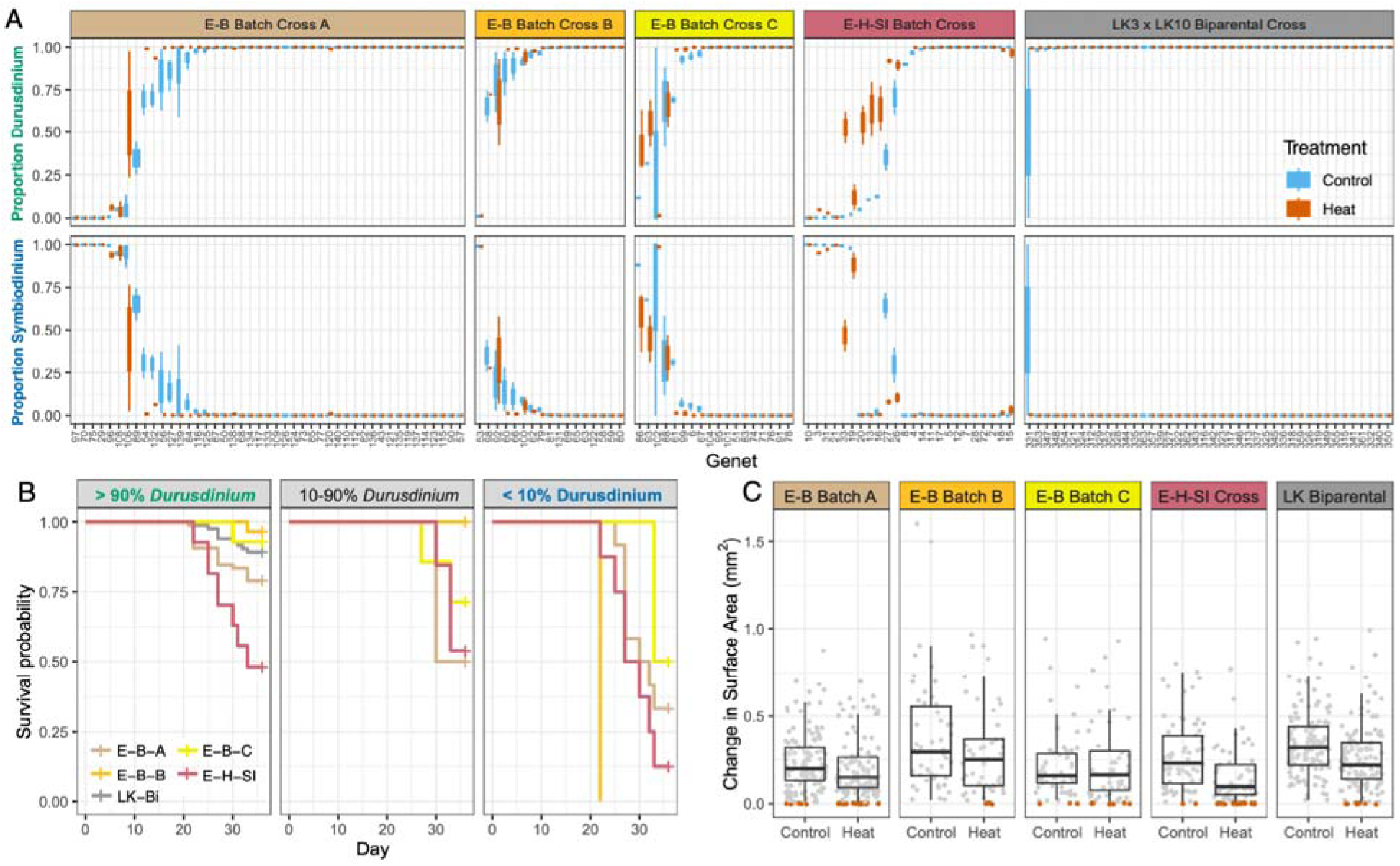
Endosymbiont community composition, growth and survival of experimental coral. (A) Proportional abundance of *Durusdinium (∼trenchii)* and *Symbiodinium (∼fitti)* within focal *Acropora palmata* genets by family origin and temperature treatment. Genet order along the x-axis is identical across panels and reflects rank order according to the mean proportion of *Durusdinium* in control samples. (B) Survival probability of heat-treated ramets by *Durusdinium* abundance and family origin. Ramets were grouped according to the relative abundance of *Durusdinium* hosted: ‘Negligible’: <10% (N=21 ramets), ‘Transitional’: 10-90% (N=27 ramets), or ‘Dominant’: >90% (N=256 ramets). Note that all LK Biparental Cross genets were classed as *Durusdinium* dominant and thus do not appear in the last two panels. (C) The change in maximal planar surface area as a function of family origin and treatment. Boxplot distributions are restricted to cases where some surface area growth was observed (∼conditional model) whereas points show all data colored by instances of no change in surface area (red) vs a positive change (grey) by cross type and treatment (∼zero-inflated model).

Symbiont density and photophysiology were also influenced by the relative abundance of *Durusdinium* hosted by a ramet in addition to family and treatment effects. Symbiont to host cell (S:H) ratios of surviving ramets were strongly influenced by the proportion of *Durusdinium* hosted (F_1,165_ = 17.0, P<0.001) as well as family origin (F_4,140.32_ = 4.16, P=0.003, Table S1, Fig. S2). Higher S:H cell ratios were observed in *Durusdinium* dominated genets, consistent with prior reports^42,43^, and pairwise post-hoc testing revealed elevated S:H cell ratios in Biparental Cross genets relative to E-B Batch Cross family A genets on average (Tukey’s HSD P = 0.01, Fig. S2). No effect of treatment or the origin by treatment interaction was detected. The proportional change in effective quantum yield (Fv/Fm) of the symbiont photosystem over the course of the experiment was also influenced by the proportion of *Durusdinium* hosted by a ramet (F_1,213.19_ = 49.1, P<0.001) as well as family origin (F_4,145.9_ = 11.9, P<0.001) and the interaction between full-sibling family and treatment (F_4,500.06_ = 5.2, P=0.0004, Table S2, Fig. S3). Genets transitioning away from *Durusdinium* exhibited greater acclimational changes in Fv/Fm over the course of the experiment (Fig. S3). Post-hoc testing revealed that the strongest treatment effects were observed in E-B Batch Cross family A genets, with heat-treated ramets maintaining higher Fv/Fm values on average than control ramets (Tukey’s HSD P = 0.004, Fig. S3), although this may be skewed by a lack of Fv/Fm data for *Symbiodinium*-dominated genets in the heat treatment. Baseline differences in control conditions were also evident, with E-B Batch Cross family A (Tukey’s HSD P<0.001) and family B genets (Tukey’s HSD P<0.05) in control conditions exhibiting greater acclimational changes in Fv/Fm than E-B Batch Cross family A, E-H-SI Cross and Looe Key Biparental cross genets.

### Relative abundance of Durisdinium explains variation in holobiont survival and partial mortality under heat stress, but not growth of surviving ramets

Complete ramet mortality was strongly impacted by treatment: of the 105 instances of ramet mortality recorded prior to and during the final sampling period, all occurred under heat treatment. Within heat-treated ramets, mortality risk over the course of the experiment was influenced by the proportion of *Durusdinium* hosted by a ramet as well as family origin (Fig. 2B). Increased relative abundances of *Durusdinium* reduced ramet mortality risk under heat treatment (hazard ratio: 0.27, P=0.0003). This effect was evident even for ramets “in transition”, defined as hosting relative abundances of *Durusdinium* between 10-90%, where mortality risk was significantly improved on average relative to ramets hosting *Durusdinium* in abundances less than 10% (hazard ratio: 2.99, P=0.01, Fig. 2B). Family effects were also evident in addition to the effects of *Durusdinium* relative abundance, and were most evident in *Durusdinium* dominated genets (Fig. 2B). The highest survival was observed in E-B Batch Cross family B, with 90% of ramets surviving to the final sampling time-point on average. Mortality risk under heat treatment was elevated in E-H-SI Batch Cross genets relative to E-B Batch Cross family B (hazard ratio: 7.994, P=0.0053), family C (hazard ratio: 4.26, P=0.0032) and the Biparental Cross (hazard ratio: 4.426, P=0.00032). Heat-treatment mortality risk was also elevated in E-B Batch Cross family A genets relative to family B (hazard ratio: 4.276, P=0.048) and Biparental Cross genets (hazard ratio: 2.37, P=0.029).

The probability of observing partial mortality was also strongly impacted by treatment, with heat treated ramets four times as likely to exhibit any degree of partial mortality as ramets in control conditions (Fig. S4, Table S3). When considering only ramets under heat treatment, both the proportion of *Durusdinium* hosted and family origin each marginally influenced the probability of observing any partial mortality, but strongly affected the extent of partial mortality when observed (Fig. S5, Table S4). Similar to the findings for complete mortality (Fig. 2B), lower partial mortality was observed on average in ramets hosting a higher proportion of *Durusdinium* (P=0.00023). Genets originating from the E-H-SI Batch Cross also exhibited higher partial mortality on average than E-B Batch Cross family B genets (Tukey’s HSD P = 0.047).

However, the proportion of *Durusdinium* did not impact the probability of observing any growth in surface area nor the increase in surface area when growth was observed (Fig. 2C, Table S5). Whereas family origin explained a significant portion of the variation in growth when observed (P<0.001, Table S6). Heat treatment increased the probability of observing no change in surface area over the course of the experiment, by 1.7 times on average relative to control conditions (Fig. 2C, P<0.0001). In cases where positive growth in surface area was observed, the effect of heat treatment on growth tended to be modulated by family origin (P=0.06, Table S6), but post-hoc testing indicates this effect was likely driven by growth differences among families in the control treatment. E-B Batch Cross family B genets exhibited greater gains in surface area in the control treatment than either E-B Batch Cross family A (Tukey’s HSD P < 0.001) or E-B Batch Cross family C genets (Fig. 2C, Tukey’s HSD P = 0.012). Growth of LK Biparental Cross genets was also higher than E-B Batch Cross family A (Tukey’s HSD P = 0.006) in control conditions and tended to be higher than E-B Batch Cross family C genets as well (Fig. 2C, Tukey’s HSD P = 0.08). Similarly, daily weight gain was influenced by family origin but not treatment nor the proportion of *Durusdinium* hosted by a ramet (Fig. S6, Table S7). E-H-SI Batch Cross genets exhibited higher daily weight gain on average than both the E-B Batch Cross family A (Tukey’s HSD P < 0.001), E-B Batch Cross family C (Tukey’s HSD P = 0.033), and LK Biparental Cross genets (Tukey’s HSD P = 0.0026).

### Genomic signatures of host propensity to retain a heterologous endosymbiont, Durusdinium trenchii, survival and surface area growth

Given the significance of endosymbiont community composition and family origin on coral holobiont traits over the course of our experiment (Fig. 2), we undertook a series of genome-wide association tests to determine if any host loci were associated with four focal traits: (1) *Durusdinium* dominance coded as a binary (≥ 90% *Durusdinium* = 1, < 90% = 0), (2) mean genet surface area growth in the control treatment, (3) genet survival under heat stress (100% ramet survival = 1, < 100% = 0), and (4) the proportional change in mean genet surface area growth in heat relative to the control treatment, after accounting for genet relatedness ^37^ and the proportion of *Durusdinum* hosted by a genet when warranted (traits 2-4).

Consistent with the strong genet-specific differences in algal endosymbiont community composition, we identified seven peaks that exceeded the genome-wide significance threshold for the genets dominated by *Durusdinium* (P < 5 x 10^-8^, Table S8, Fig. 3B,C,S7). Gene density varied across the three regions with consistent associations, with most containing between 14-27 genes within 200–300 kb windows, of which between 3-8 were annotated (Table S9). Annotated genes located in proximity to these three peaks, included MC4R (2 copies), MC5R, CD163, EIF3G, WDR83, C9orf114, CERCAM, MAFG (2 copies), MAF, ALG6, RFX3, ERBB2IP, CKMT2 (2 copies), NDC1, and EMC1 (Table S9). The chromosome 1 peak was the only region where SuSiE fine-mapping resolved a single credible set (PIP > 0.95), encompassing 14 tightly linked SNPs (Fig. 3A, Table S10). Although no annotated genes directly overlapped with the putative causal sites identified in this credible set, the region includes two copies of a gene annotated as a melanocortin 4 receptor (MC4R) out of only seven copies in the entire genome (Fig. 3A). Functional enrichment analyses of annotated genes near this peak further emphasized this finding, with significant enrichment of melanocortin receptor activity (GO:0004977) and melanocyte-stimulating hormone receptor activity (GO:0004980) among molecular function (MF) terms (Table S11). It is important to note, however, that these annotations should not be used to infer function. Genes annotated as MC4R were detected in multiple anthozoan genomes (Blastp e-value < 10^-20^) but no endosymbiont genomes indicating host-specific origin and evolutionary conservation (Table S12). Yet homology was low, with only around 30% identity to chordate melanocortin receptors on average (Fig. S8). Nor could we find homologs for melanocortin receptor ligands (AgRP, MSH) or their precursors (ASIP, POMC) in any coral or endosymbiont genome (Table S12).

**Figure 3.**
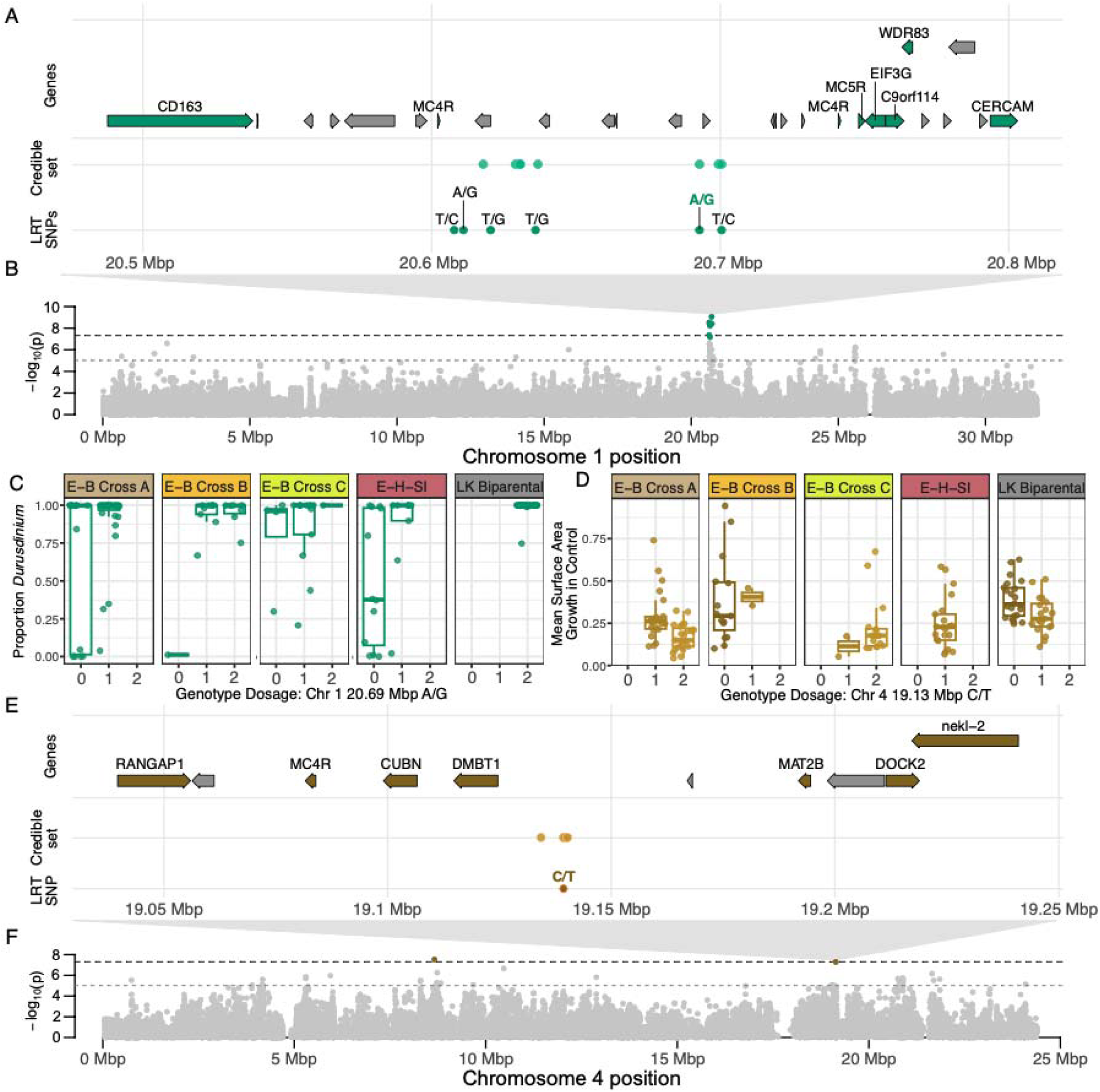
Genomic regions associated with coral host propensity to retain *Durusdinium* and mean surface area growth in control conditions. (A) Relative position of significant SNPs within focal peak for propensity to retain *Durusdinium*, the associated credible set resolved during fine mapping, and genes in proximity. Unannotated genes are shown in grey. (B) Manhattan plot of P-values for SNPs on Chromosome 1 with the propensity to retain *Durusdinium,* coded as a binary (≥ 90% =1,< 90% =0), obtained from the genome-wide association test. (C) Boxplot distribution of actual *Durusdinium* proportion by genotype dosage (0: homozygote for reference allele, 1: heterozygote, 2: homozygous alternate) at most significant SNP by family origin. (D) Boxplot distribution of mean surface area growth in control conditions by genotype dosage at focal SNP by family origin. (E) Relative position of significant SNP associated with surface area growth in control conditions, the associated credible set resolved during fine mapping, and genes in proximity. Unannotated genes are shown in grey. (F) Manhattan plot of P-values for SNPs on Chromosome 4 with surface area growth in control conditions obtained from genome-wide association test. Horizontal dashed lines on Manhattan plots indicate genome-wide suggestive (grey, P < 1 x 10^-5^) and significance threshold (black, P < 5 x 10^-8^).

Given the minimal influence of endosymbiont type on surface area growth (Table S5,S6), we also explored whether any regions in the genome were associated with surface area growth in control conditions. We identified one peak that exceeded the genome-wide significance threshold (P < 5 x 10^-8^) and one which almost met the threshold (P = 5.24 x 10^-8^) both of which were localized to chromosome 4 (Fig. 3F,S9). The first peak on chromosome 4 contained nine genes, of which seven were annotated as GRN, CHM, SLC25A36 (2 copies), TAF1, ALKBH8, and KCTD1(Table S9). The second peak contained 10 genes, with seven annotated, including RANGAP1, DMBT1, MAT2B, CUBN, MC4R, DOCK2 and nekl-2 (Table S9). SuSiE fine-mapping resolved a 95% credible set in this second peak comprising 11 tightly linked SNPs of which 5 exhibited posterior probabilities > 0.05 (Fig 3E, Table S10). The lead SNP in the set is located only ∼9.7 kb downstream of the 3’ end of DMBT1 and functional enrichment analyses generally supported the involvement of pathways regulating growth and immune responses. Biological process (BP) terms enriched near the first peak for functions related to mononuclear cell proliferation (GO:0032943;GO:0070661;GO:0046651) and T cell proliferation (GO:0042098; Table S11).

As ramet mortality in heat was influenced by batch cross family origin in addition to symbiont community composition, we also explored whether any regions in the genome were associated with the probability of 100% survival in the heat treatment. While four regions exceeded the genome-wide suggestive threshold, none exhibited strong associations with the proportion of ramet mortality (Fig. S10). This is aligned with prior studies in which the genomic basis of coral thermal tolerance appears to be highly polygenic^5,9^ and thus our sample size is likely not sufficient to detect loci of small effect. Similar to percent mortality, no regions exceeded the genome-wide significance threshold for associations between proportional change in surface area growth under heat treatment relative to control, but seven exceeded the genome-wide suggestive threshold, with three showing reasonably consistent relationships between phenotype and genotype dosage: two localized to chromosome 4 (bp13288405 P = 2.6 x 10^-6^, bp20154249 P = 4.1 x 10^-7^) while the third localized to chromosome 8 (P = 2.4 x 10^-6^, Fig. S11, Table S8). Genes in proximity to these loci included ACIN1, RPL9, CLCN3, CRAMP1L, TRM10, TXNDC9, GLYCTK, DAG1, FAM208A, C1orf101, NDUFAF7, and MGAT4C (Table S9).

### Differential expression of MC4R is associated with changes in symbiont community composition following transplantation to natural reefs

Given the strong associations detected for symbiont community composition and surface area growth in control conditions (Fig. 3), we leveraged samples from an independent experiment^39^ to quantify host gene expression associated with changes in symbiont community composition and surface area growth following transplantation of five *A. palmata* genets to natural reef sites in the Lower Florida Keys (Fig. 1D-F). Genet identity did not explain the presence of *Symbiodinium fitti* (p = 0.908) or changes in dominant symbiont type (p = 0.989) after field transplantation^39^ (Table S13) although there was a trend of chromosome 1 peak heterozygote genets retaining more *Durusdinium* at T24 (Fig. S12), consistent with the GWAS signal (Fig. 3C). Differential expression analysis identified two genes that were significantly differentially expressed (|log2FC| > 1 and P_adj_ < 0.05) in outplanted ramets in which any *Symbiodinium* was detected and in which *Symbiodinium* became the dominant community member (Fig. 1G). In both contrasts, g18865 (TBCD) and g7717 (MC4R) were upregulated in samples dominated by *Symbiodinium fitti* (Fig. 1F,4C,4D; Table S14). No additional genes met the significance threshold. This copy of MC4R is located on chromosome 4, in proximity to the GWAS peak associated with surface area growth (Fig. 3E). However, there was no association between genotype dosage at the outlier SNP in that region and symbiont community composition (Fig. S13).

### RANGAP1, a gene in proximity to growth-associated SNP, is upregulated in coral transplants with the highest surface area growth on natural reefs

To explore correlations between gene expression and surface area growth two-years post transplantation to natural reef sites, expression was modeled as a function of relative surface area growth using linear models with and without transplant destination (Fig. 1A,E) and dominant symbiont type (Fig. 1F) as covariates. Ramet surface area increased over time (F_4,28_ = 275, p < 0.001; Fig. 1E) and was also influenced by the additive effect of outplant site (F_7,28_ = 3.03, p = 0.032, Fig. 1E). Surface area growth was greatest at Eastern Dry Rocks (Fig. 1E), but pairwise differences among sites were not significant after correcting for multiple testing. A total of 19,174 genes were retained after low-expression filtering. Of these, 49 were significantly differentially expressed as a function of relative surface area growth (Fig. 4A). All significant associations were detected in the univariate model (expression ∼ relative SA growth), and inclusion of location or dominant symbiont as covariates did not yield additional significant genes. Of these 49, 41 were upregulated and 8 were downregulated among high growth ramets (Fig. 4A), and 15 had gene name annotations (Table S15). No significant functional enrichments were detected, but gene names are suggestive of roles in morphogenesis and proliferative growth (TGFB3), extracellular matrix formation (HMCN1), and stress-response and proteostasis (PRDX4). We also investigated overlap between these 49 DEGs and the 19 genes in proximity to the two outlier regions on chromosome 4 associated with surface area growth in control conditions (Fig. 3E). One significantly differentially expressed gene, Ran GTPase-Activating Protein 1, is single-copy and located in proximity to the second peak on chromosome 4 and was significantly upregulated in ramets with the highest surface area growth on natural reef sites (RANGAP1, g7715, Fig. 3E,4B Table S15).

**Figure 4.**
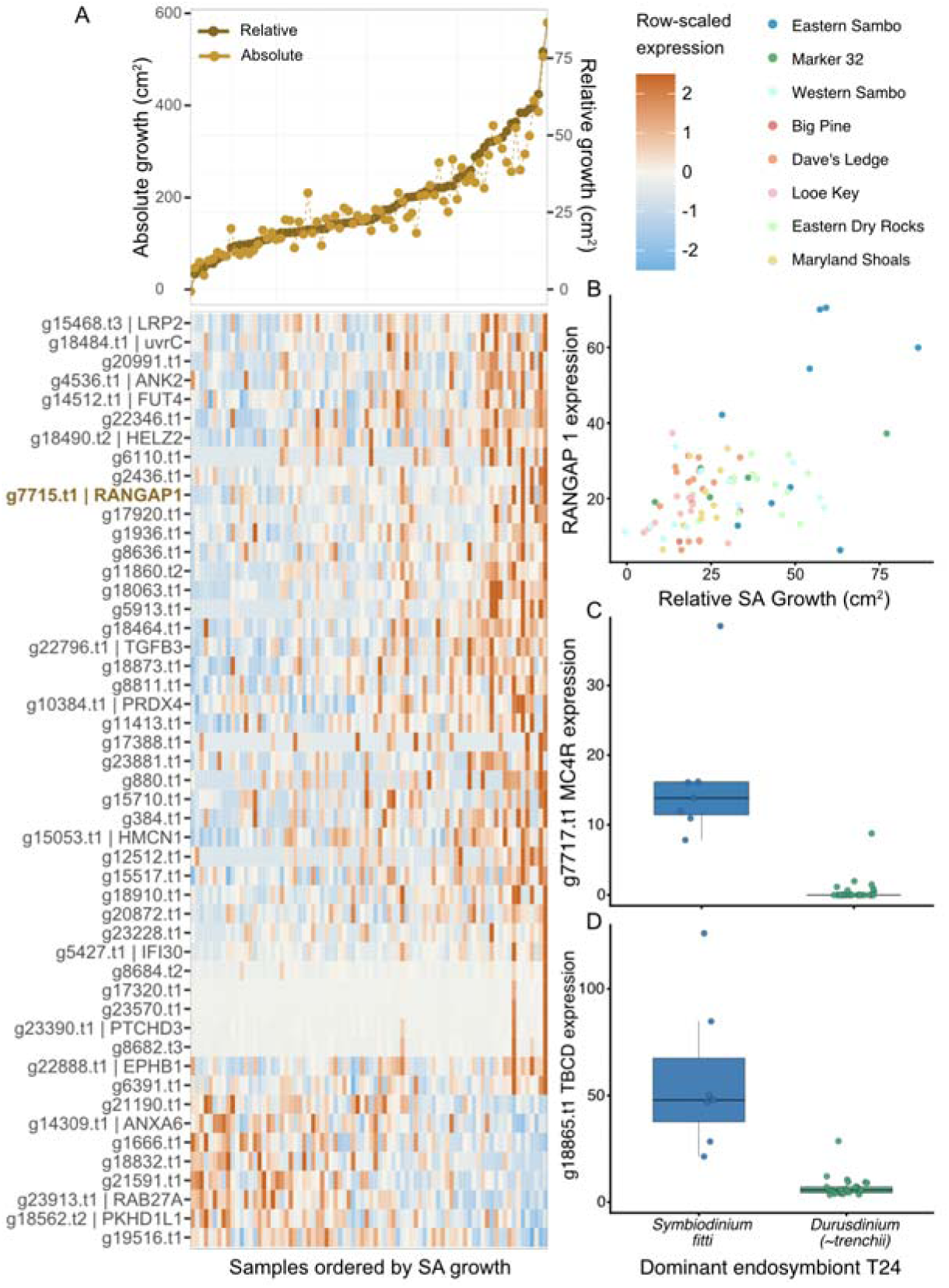
Global gene expression patterns in *A. palmata* associated with symbiont shuffling and surface area growth on natural reefs. (A) Heatmap showing row-scaled expression of 49 differentially expressed genes significantly associated with relative surface area growth. Samples (∼columns) are ordered according to their relative surface area (SA) growth as shown in line plot inset above heatmap. (B) Absolute expression of RANGAP 1, a gene in proximity to the credible set on Chromosome 4 associated with surface area growth in the controlled tank experiment, by relative surface area growth in field transplants colored by site of outplant. Absolute expression of (C) MC4R (g7717.t1) and (D) TBCD (g18865.t1), the two genes significantly differentially expressed in coral ramets by their dominant endosymbiont type, two-years post-transplantation to natural reef sites in the Lower Florida Keys.

## Discussion

Using common-garden reared genets of *Acropora palmata*, we identified genomic regions associated with areal growth and endosymbiont community composition which also exhibited a functional role in coral transplanted to natural reefs. Specifically, two copies of a gene annotated as a melanocortin 4 receptor (MC4R) and one annotated as melanocortin receptor 5 (MC5R) were located in proximity to a strong outlier peak on chromosome 1 associated with propensity of coral genets to retain *Durusdinium* in the common garden coral nursery, while a separate copy of a putative MC4R was upregulated in ramets which naturally shuffled to *Symbiodinium fitti* in the field. More strikingly, expression of RANGAP1, a Ran GTPase activating protein, was positively correlated with surface area growth in coral outplants and was located 78.3kbp upstream of an outlier peak associated with areal growth of coral genets in the common garden. While these independent lines of evidence provide strong support for the functional significance of these candidate genes and contribute to our understanding of both how phenotypic variation evolves and how it can be harnessed for conservation applications, there is a need to further investigate the basic biology of these associations. RANGAP1 is a conserved marker of cell division, but its functional significance has only been delineated in laboratory models to date. Similarly, the melanocortin system is thought to be restricted to vertebrates, yet our results suggest homologs could play similar functional roles in basal anthozoans. Elucidating the mechanistic roles of these genes in corals will therefore be critical for understanding their adaptive significance and for translating genomic discoveries into effective reef restoration strategies.

### Ran GTPase-Activating Protein 1 is a conserved marker of cell division

RANGAP1 is the GTPase-activating protein for Ran, a subfamily of the Ras GTPase superfamily. Activation of Ras superfamily members by extracellular stimuli triggers signalling cascades involved in activation of fundamental cellular processes, including cell growth and differentiation. The Ras GTPases have been implicated in the coral stress response^44^ and Rhohomologs were identified in all basal metazoa^45^ but the Ran family, which includes RANGAP1, has not been a major focus of work in coral or related cnidarians (e.g. *Hydra vulgaris*, *Nematostella vectensis*). However, its role in mitotic cell division as a key regulator of the Ran GTP/GDP cycle is well-established in other model systems^46^. The RanGTP cycle plays a significant role in nuclear transport, with RANGAP1 involved in cytoplasmic localization^47^. Its role appears evolutionarily conserved as a homolog in *Arabidopsis* is a protein marker of the plant cell division plane, dependent on known regulators of plant cytokinesis^48^ and in the yeast, *Schizosaccharomyces pombe*, it contributes to heterochromatin formation through associations with both histone H3 and the H3-lysine 9 methyltransferase^49^. Interestingly, RANGAP1 was recently shown to regulate bone development in mice^50^, thus it is tempting to speculate that it plays a similar role in coral skeletal formation. As mitotic cell division is the foundation of somatic growth, an alternative or additional role for RANGAP1 may be in regulating soft tissue growth. The positive correlation we detected between RANGAP1 expression as well as proximal genetic variation and surface area growth aligns with either hypothesis, although the mechanism of coral calcification is different from mammalian osteogenesis^51^. Additional functional work, such as localization of its expression and/or protein product, will be necessary to verify its role in coral.

### Prospective role of melanocortin-like receptors in the coral symbiosis

The melanocortin 4 receptor is a major coordinator of energy homeostasis and body weight in mammals^52^. Its role has been best characterized in neurons, where neuropeptides AgRP (Agouti-Related Peptide) and α-, β- and γ-melanocyte stimulating hormone (MSH), inhibit or enhance neuronal signalling, respectively^52–54^. Importantly, MC4R has a well-established role in central nervous system signalling and is distinct from peripheral melanocortin receptors involved in pigmentation, such as MC1R^55^. Pharmacological manipulations have shown that MC4R agonists (or ligand analogs which activate receptor signalling) promote satiety, energy expenditure and weight loss whereas antagonists (compounds which inhibit receptor function) increase food intake, energy conservation and weight gain^53,54^. Given that the coral symbiosis is fundamentally nutritional, and that there is variation among symbiont species in host nutritional provisioning with *Durusdinium* spp. known to translocate less carbon^18,22^, it is tempting to speculate that MC4R plays a similar role in coral, sensing and responding to the availability of symbiont-derived carbon, which could influence propensity to associate with specific partners.

However, the melanocortin system evolved in vertebrates and while homologs have been identified in protochordates, even these do not recognize the canonical ligands^55–57^. Moreover, echinoderms, cephalochordates, urochordates and protostomes lack melanocortin receptor homologs in their genome^58^, although Pomc-like products that encode one or more melanotropin-like peptides were isolated from the bivalve, *Mytilus edulis*^59^ and the leech, *Theromyzon tessulatum*^60^. Surprisingly, we identified 22 genes annotated as melanocortin receptors in the *A. palmata* genome, of which seven were annotated as MC4R using emapper^61^. Melanocortin gene annotations were consistent features of multiple anthozoan genomes indicating some degree of evolutionary conservation, yet the overall percent homology is low. Nor could we find putative homologs for the ligands (AgRP, MSH) or the MSH precursor (proopiomelanocortin/POMC) in any coral or endosymbiont genome. However, MC4R is known to exhibit significant basal activity in the absence of signalling ligands, and thus agonists may not be necessary for function^55^. Future work should prioritize comparative genomic investigations of these melanocortin receptor annotations in anthozoan and other related genomes, including cellular localization of MC4R and identification of ligands to better understand its role in the coral symbiosis.

### A potential role of distal enhancers in coral phenotypic variation

While there is substantial support for the role of cis-regulatory variation in phenotypic evolution^38^, the role of distal enhancers has only been recently established through the advent of genomic tools capable of interrogating chromosome topology^62^. The exact mechanism has yet to be resolved, but one model gaining traction posits dynamic turnover of transcription hubs where enhancers need only need be in proximity to target promoters to alter transcription, with a distance of 100-300 nm established as sufficient in *Drosophila* embryos^62,63^. Work in model systems has shown that enhancers can simultaneously activate multiple promoters linked in cis (on the same chromosome) or in trans (on different chromosomes)^63,64^, while coordination among enhancers on separate chromosomes can yield activation of single genes^65–67^. We identified genomic variants correlated with the propensity to retain *Durusdinium trenchii* during shuffling on chromosomes 1,3 and 14, but none occurred within coding regions. Annotated genes near the peak on chromosome 1 were enriched for melanocortin receptor activity, while the copy of MC4R that was differentially expressed in coral which exhibited shuffling in a field transplant was located on chromosome 4. We therefore speculate that a similar distal enhancer mechanism mediated through structural changes in the relative proximity of chromosomes, as observed in *Drosophila*, could explain this pattern. Specifically, we hypothesize the peak on chromosome 1 could be in proximity to an enhancer for the two cis MC4R promoters as well as the trans MC4R promoter on chromosome 4. Similarly, the SNP variants on chromosome 4 associated with differential surface area growth are proximal to, but do not fall within coding regions. We again speculate that this region may serve as a cis-enhancer for our top candidate gene, RANGAP1. At present, however, the role of enhancers and chromosome topology in regulating expression in non-model systems can only be inferred. Annotating coding regions remains a major problem in non-model organisms, including coral^44^, and the functional role of non-coding variation remains almost entirely unknown. Future work, such as chromatin immunoprecipitation-type approaches^68^, may provide more insight into prospective long-distance interactions.

### Application to coral conservation and restoration

From an applied interventions perspective, some of our findings are immediately actionable while others represent targets for continued research and development. First, our results suggest that manipulation of endosymbiont associations can increase coral fitness with minimal cost. Similar to the findings of Turnham et al. (2023)^69^, association of *A. palmata* with *D. trenchii* did not incur a growth trade-off in our experiment: LK Biparental Cross genets, which were nearly all *Durusdinium* dominated exhibited some of the highest areal growth on average. Moreover, transitional and *Durusdinium*-dominated genets also exhibited higher mean survival in the heat treatment relative to *Symbiodinium*-dominated genets. Enhanced survival could help offset early-life mortality in support of higher restoration yields. In addition, an initial association with *D. trenchii* did not limit the capacity of these genets to associate with the homologous symbiont, *Symbiodinium fitti*, as all genets in the nursery ultimately shuffled to *S. fitti* dominance (E. Muller, pers. communication) and genets were capable of shuffling in the field^39,40^. This suggests that even if growth deficits were to manifest, they would not be irreversible. Taken together, these findings can inform risk-benefit analysis by regulators considering restoration intervention proposals involving heterologous symbiont association ^70^. However, it is important to note that the effect of *D. trenchii* may be partially rescued by other host genetic variation favorable for growth. LK Biparental genets comprise genotypes most positively correlated with surface area growth, which extends reports of species-level interactions between host and symbiont^71^ to variation among individuals and provides a mechanistic explanation for this phenomenon.

Building on this, we also find a role of host genetic variation in structuring endosymbiont associations, survival under heat stress, and areal growth in support of host genetic interventions. Specifically, we identify genotypes which are more likely to retain *Durusdinium trenchii* even under thermal stress, genotypes exhibiting higher growth on average, and coral families exhibiting higher survival that is not attributable to their dominant endosymbiont type. Moreover, although our sample size is small, we show that the direction of these relationships remains consistent outside of a controlled lab environment and genes implicated in producing these phenotypes in a controlled laboratory setting are also predictive of trait values in the field. Specifically, the expression of an MC4R gene was associated with endosymbiont type and RANGAP1 expression was positively correlated with areal growth over two years in the field. This indicates that approaches such as selective breeding and assisted gene flow could be used to increase the frequency of these alleles and their associated traits in restoration populations^72^, and that biomarkers could be developed to predict phenotype^2^. Encouragingly our finding that specific families of *A. palmata* can exhibit both higher survival under heat stress and higher growth while associating with *D. trenchii* also aligns with the emerging body of work reporting a general absence of trait trade-offs in other coral species^69,73–75^. Again, these findings support the purported benefits of host genetic interventions and indicate that risks remain minimal, assuming that broad genetic variation is appropriately managed^76^.

As restoration programs incorporate experimental bleaching stress as a screening method for climate-smart interventions, it is imperative to determine whether the exposures and trait metrics used for ascribing tolerance are useful as selective tools. Interestingly, we did not observe an effect of heat treatment on S:H ratios and Fv/Fm, yet mortality was observed under the month-long thermal exposure. Fv/Fm is an increasingly used metric in screening genotypes to identify enhanced tolerance to improve restoration outcomes^77–80^. However, we found an increased proportional change of Fv/Fm in heated ramets with greater proportions of *D. trenchii*, suggesting photochemistry in this species may not be affected under bleaching stress (see also^78^) and supporting greater thermal tolerance in corals associating with *Durusdinium* spp.^16,19,69,81^. Although ramets experienced mortality under thermal stress, *in hospite* symbionts are still photophysiologically functional; therefore, Fv/Fm may not always be a reliable indicator of visible coral bleaching and its subsequent impacts^82,83^.

Finally, our results also underscore the need to better understand non-coding variation in driving coral phenotypes. The relevance of cis-regulatory variation in driving phenotypic evolution has been well-established for decades^38^, but its role in conservation and restoration contexts is underappreciated. For example, gene editing has been heralded as one means of accelerating coral evolution in the face of climate change^44,72^, but to date studies have focused on perturbing coding sequences^84–86^. Epigenetic modifications have been proposed^87,88^ in addition to transgenics targeting over/underexpression of particular candidate genes^89^, which could represent a viable alternative to direct modification of non-coding regions, but the technologies have yet to be tested in coral.

### Limitations and Considerations for Future Work

The limited genetic diversity of our study population constrained the suite of analyses we could undertake, necessitating further validation studies. Our sample population comprised several full-sibling groups^37^ and high relatedness precludes calculation of polygenic risk scores due to violation of the assumption of independence among individuals^90^. Moreover, while we did not identify any thermal tolerance associated loci, our phenotypes were limited. Since our Fv/Fm and S:H cell ratio metrics did not respond strongly to a month-long thermal challenge, possibly due to the photophysiological nature of *D. trenchii*^78^, we were not confident in these phenotypes bearing strong merit for a GWAS, and instead focused on mortality and relative growth deficits. This reinforced our decision to incorporate additional evidence from an independent field transplant experiment^39^ to interrogate candidate gene function, but it should be noted that a broader regional sample set could uncover other important variants. That being said, while *A. palmata* populations exhibit regional substructure, average genomic divergence is relatively low^91^ suggesting that the allelic variants we identified are likely to be regionally widespread.

Finally, even if heritable genetic variants can be identified and leveraged in restoration applications, extreme environmental conditions may still overwhelm a genetic predisposition for a particular phenotype. The magnitude of thermal stress experienced on Florida reefs in 2023/24, nearly three times that ever previously recorded, underscores this threat: 98% of Floridian acroporids died regardless of their trait values or genotypes^8^. Thus it will be essential for future work to establish the ultimate utility of harnessing this genetic variation for future restoration applications given the rapid rate of climate change. At a minimum, there is utility in biobanking genomic variation and thus our findings could inform selection of individuals to incorporate into these efforts.

## METHODS

### Moderate duration thermal stress experiment

*A. palmata* coral were sourced from Mote Marine Laboratory’s *ex-situ* coral nursery at the Elizabeth Moore International Center for Coral Reef Research and Restoration (IC2R3) facility in Summerland Key, FL (24.6617° N, -81.4554° W) on March 2, 2022. A total of 156 genotypes of *A. palmata* were fragmented into seven ∼2 x 2 cm replicate ramets (six experimental, 1 spare) and adhered onto individual ceramic plugs. The majority of these 156 genotypes were the product of sexual reproduction among founder genets from 2013-2020^37^. Specifically, analysis of kinship coefficients yielded assignment of the majority of these individuals to five full-sibling groups: 24 full-siblings derived from a 2013 batch cross between Elbow, Horseshoe, and Sand Island founder genets (hereafter the E-H-SI Batch Cross), 42 full-siblings derived from a 2020 biparental cross between two Lower Keys founder genets (hereafter the LK-3 x LK-10 Biparental Cross), and 82 that were the product of 2017 batch crosses between Elbow Reef and Brewster/Ball Buoy Reef (Biscayne National Park) founder genets which comprised three full sibling groups, hereafter referred to as E-B Batch Cross A (n=45), E-B Batch Cross B (n=17) and E-B Batch Cross C (n=18) while two genets were unrelated. Remaining genets were derived from a 2014 batch cross comprised solely of Elbow Reef parents (n=3), a 2015 batch cross between Elbow and Horseshoe reef parents (n=1), and finally, four were founder genets sourced directly from Looe Key, Sand Key, Western Dry Rocks, and Turtle Reef in 2018, 2021, 2021, and 2014 respectively (Fig. 1A,B).

After fragmentation, ramets were transferred into the IC2R3 Climate and Acidification Ocean Simulator (CAOS) system where they recovered in ambient conditions (∼27.5°C) until the start of the experiment (approximately one month post transfer). Ramets were placed into 6 raceways (3 high temperature and 3 control temperature) and randomly distributed onto 10 plastic egg crates per raceway, with 1 ramet per genotype in each raceway. Seawater was pumped into each raceway at 6.25 ml/s. On April 1, 2022, incremental increases in temperature began in the three high temperature raceways at a rate of +0.5°C/day until ∼31.5°C was reached on April 10, 2022, and temperature was maintained until the end of the experiment (Fig. 1C). Control raceways were held at ambient temperature, ∼27.5°C, throughout the experiment. The ambient temperature of ∼27.5°C reflected the average ambient sea water temperature between March and May in the Florida Keys and the elevated temperature, ∼31.5°C, reflects +1°C above the bleaching threshold. Water quality was recorded on a daily basis using YSI handheld and pH probe (Mettler Toledo). Each raceway was fed supplementary nutrition twice a week with a mixture of Microvore (Brightwell Aquatics, Fort Payne, AL, USA), Zooplankos-S (Brightwell Aquatics, Fort Payne, AL, USA), and Reef Snow (Brightwell Aquatics, Fort Payne, AL, USA). Husbandry occurred on a daily basis and involved the removal of algae and detritus from each raceway and individual ceramic plugs. Systems were covered with shade cloth from 11AM-3PM to minimize UV stress.

### Phenotypic trait measures

Measurements of buoyant weight, coral skeletal/tissue surface area and photochemical efficiency were performed prior to (T_initial_ = 0 days) and after (T_final_ = 33-36 days) exposure to heat stress. Ramet survival was tracked during the course of the experiment and any ramet exhibiting more than 30% tissue loss was removed and the time of death was recorded resulting in a tally of alive (1) and dead (0) ramets at the end of the experiment, which was at 36 days for control corals and 33 days for heat treated corals. At the end of the experiment and after the conclusion of all measurements, living corals were wrapped in foil and flash frozen for subsequent nucleic acid extraction.

At both time points, all ramets were photographed using an Olympus (TG-6) that was mounted to an overhead rig. Ramets were oriented in a fixed position with a ruler in the frame for scale and a coral watch color health chart in the frame for color comparison. Ramet skeletal surface area (mm²) was measured from photographs by tracing the entire fragment using TagLab™ (Pavoni et al. 2022) following the methods outlined in (Amir et al. 2023). Ramet live tissue surface area (mm²) was measured from photographs by tracing only the edges of the live tissue on each fragment using TagLab. Maximal surface area growth was calculated as the difference between skeletal surface area at the final time-point and the initial surface area of experimental ramets. Partial mortality was calculated as the difference between skeletal surface area and live tissue area at the final time-point.

Ramets were dark adapted for 30 minutes before measurements of Photosystem II (PSII) photosynthetic efficiency were taken using an Imaging PAM (Walz). The area of interest (AOI) was positioned at the center of each fragment, in an area with no tissue recession, before each measurement was collected. Settings used were measuring light intensity = 6, SST-Pulse Intensity = 12, SAT-Pulse width = 0.6, and gain = 2. Fv/Fm (the maximum potential quantum efficiency of PSII), F_0_ (the fluorescent light in the absence of actinic photosynthetic light), NPQ (the efficiency that light is converted to heat), ETR-Max (maximum electron transport rate (µequiv m ²s ¹)) (Maxwell and Johnson 2000) were collected from fluorescence measurements. Proportional differences in aspects of symbiont photophysiology (Fv/Fm, NPQ and ETR-Max) were calculated as the difference between final and initial values standardized to the initial value.

The buoyant weight of corals were measured following the protocol outlined in Davies (1989), where corals were placed on a submerged weighted platform suspended under a precision balance on 10-gallon aquarium tanks containing 35 L of seawater from respective treatments. Temperature and salinity were measured hourly to calculate the density of seawater within each of the weighing tanks. One coral replicate per rack was weighed twice to determine balance accuracy (0.217 ± 0.489 g). Dry weight was calculated following the methods in Jokiel et al. 1979. As coral were buoyant weighed on consecutive days, we calculated Daily Weight Gain as the difference in air converted weights between the final and initial time-points standardized by the number of days between consecutive weighings.

### S:H cell ratios and symbiont community composition

Holobiont DNA was extracted from a representative ramet of each genet using a modified CTAB protocol^92^. Twelve samples failed to yield sufficient DNA and were excluded from further analysis of symbiont to host (S:H) cell ratio and community composition analysis. Relative abundance of the three Symbiodiniaceae genera previously shown to associate with captive bred *A. palmata* (*Symbiodinium*, *Cladocopium*, and *Durusdinium*^39,40^) and coral host cells were quantified with 20 ng input DNA using actin-based qPCR assays on an AriaMx real-time PCR system (Agilent Technologies, USA) following^43,93^. A total of three qPCR reactions were run per sample, each with two technical replicates. One TaqMan (Thermo Fisher Scientific) reaction targeted the *Symbiodinium* (formerly Clade A) actin locus with final concentrations of 300 nM probe, 300 nM Aact forward primer and 200 nM Aact reverse primer in a 25 ul total reaction volume. A separate TaqMan (Thermo Fisher Scientific) multiplex reaction in a 25 ul final reaction volume targeted the actin locus in *Cladocopium* (formerly Clade C) and *Durusdinium* (formerly Clade D) with final reaction concentrations of 100 nM per probe, 50nM forward primer and 75 nM reverse primer^43^. A final reaction amplified the host calmodulin locus with SYBR-Green again in a 25 ul volume with final concentrations of 200 nM for each CaM primer^94^. An additional melt curve step was run for the host reaction to verify a single template indicating specificity. Raw cycle threshold (CT) values were corrected for differences in fluorescence intensity between the reaction-specific fluorophores, determined by creating standard curves of target amplicons ranging from 10^8^-10^2^ which revealed that the *Cladocopium* assay produced CT values 1.23 cycles higher than *Durusdinium* and *Symbiodinium* and that the SYBR-Green assay amplified 6.3 cycles sooner than the *Symbiodinium* probe and 5.3 cycles higher than the *Cladocopium* and *Durusdinium* probes on average (Fig. S14). S:H cell ratios were then calculated from adjusted CT values using the formula from ^43^ with target locus copy number ratios based on ^94^, Table S16).

### QC, visualization and statistical analysis of traits

All data summaries, figures and statistical analyses were undertaken in R v4.4.2^95^. Raw trait data were inspected and filtered for erroneous trait measures. Skeletal surface area measures for one ramet, 117.5, revealed a final skeletal area smaller than the initial measure (Table S17). Re-inspection of photographs did not reveal any obvious breakage and independent re-measuring of the surface area in TagLab did not change values, thus we excluded this sample from the final maximal surface area growth dataset as the discrepancy may be attributable to a change in image angle warping the size standard.

Summary plots of the buoyant weight data revealed a largely even distribution around a mean of +0.0045 g/day, but some extreme outliers (those showing an order of magnitude higher or lower growth rate) were detected in clusters. Further investigation into the metadata revealed that several extreme high and low outliers were temporally clustered in the final buoyant weight datasheet. For example, on 7 May 2022, both an extreme high (0.01 g/day) and extreme low (- 0.05 g/day) value sample were weighted at 2:50 pm. To be conservative, we elected to exclude temporally overlapping daily growth rate measures exceeding the interquartile range criterion, IQR. Where IQR is defined as values exceeding 1.5 times the IQR, or difference between the first and third quartile, below the first and above the third quartile, respectively. A total of 45 measures were identified as exceeding the IQR criterion and of these, 36 also exhibited temporal overlap and were filtered from the dataset prior to statistical analysis and plotting.

qPCR data were filtered to remove any samples with a coefficient of variation of technical replicate CT values exceeding 20% and instances where host CT values were too high to yield a confident estimate of the S:H cell ratio, defined as a mean host CT value greater than or equal to 28 cycles.

To test for the effect of family origin and treatment conditions on trait values, we restricted the quality filtered dataset to include only families with sufficient replication (>3 genets per family), resulting in five groups of full-sibling genets, three originating from Elbow Reef and Biscayne National Park founder genets (hereafter the E-B Batch Cross families A, B and C), those originating from a 2013 batch cross between Elbow, Horseshoe, and Sand Island founder genets (hereafter the E-H-SI Batch Cross), and those originating from a 2020 biparental cross between two Lower Keys founder genets (hereafter the LK-3 x LK-10 Biparental Cross^37^).

Most ramets remained dominated by *Durusdinium* (∼*trenchii*) as has previously been reported for *ex situ* reared *A. palmata*^39,40^, although some had transitioned to *Symbiodinium* (∼*fitti*) dominated communities with a small minority dominated by *Cladocopium* spp (N=3; Fig. S1). The proportions of *Durusdinium* and *Symbiodinium* were therefore modeled as a function of family origin, treatment, and their interaction using beta regression models as implemented in the betareg package^96^. A second set of models exploring genet nested within family origin and treatment was also implemented to evaluate the explanatory power of genet. Post-hoc Tukey tests were implemented with the package emmeans^97^ to evaluate the significance of pairwise contrasts among genets nested within family. Given the well-established physiological consequences resulting from variation in Symbiodiniaceae community composition, all subsequent trait analyses included the proportion of *Durusdinium* hosted by ramets as a covariate.

Maximal surface area growth was modeled as a function of the proportion of *Durusdinium*, as well as family origin, treatment, and their interaction using generalized linear mixed models with a zero-inflated Gamma family distribution (additive effects only) as implemented in the glmmTMB package^98^. Partial mortality was initially modeled similarly to surface area growth, but complete separation prevented modeling a more complex interaction as no instances of partial mortality were observed in the Biparental Cross in control conditions (Fig. S4). To more reliably estimate the effects of symbiont community composition and family origin on partial mortality the dataset was restricted to exclude control samples and the model was rerun omitting the treatment factor. Raceway was included as a scalar random effect for all surface area and partial mortality models. A Wald Chi^2^ test was used to evaluate the significance of conditional fixed effects, and when a significant effect of family or the interaction term was detected, a post-hoc Tukey test was implemented with the package multcomp^99^ to evaluate the significance of pairwise contrasts.

S:H cell ratios were log10 transformed to satisfy assumptions of linearity and heteroscedasticity and modeled as a function of the fixed effects of the proportion of *Durusdinium* hosted by a ramet, as well as family origin, treatment and their interaction. Scalar random effects of genet and raceway were also included. Significance of fixed effects was evaluated using Kenward-Roger’s method.

Daily weight gain was modeled as a function of the fixed effects of the proportion of *Durusdinium* hosted by a ramet as well as family origin, treatment and their interaction. A scalar random effect of raceway was also included, but genet was excluded to reduce effects of overfitting. Significance of fixed effects was evaluated using Kenward-Roger’s method.

The proportional change in Fv/Fm over the course of the experiment was modeled as a function of the fixed effects of the proportion of *Durusdinium*, as well as family origin, treatment and their interaction. Scalar random effects of genet and raceway were also included. Significance of fixed effects was evaluated using Kenward-Roger’s method.

A mixed-effects cox proportional hazards model was used to evaluate complete ramet mortality in the heat treatment prior to the end of the experiment as a function of the fixed effect of family origin and proportion of *Durusdinium*, including a scalar random effect of raceway. The effect of treatment could not be modeled explicitly due to a lack of variance in the control treatment (100% ramet survival) leading to an unrealistically high mortality risk coefficient and models including an interaction term did not converge. Therefore we modeled mortality risk in heat-treated ramets as a function of family origin and the proportion of *Durusdinium* hosted. The proportion of *Durusdinium* was modeled both as a continuous trait and as a categorical. For the categorical assignment, ramets with *Durusdinium* abundances of 90% or greater were considered *Durusdinium* ‘dominant’, those with *Durusdinium* abundances between 10-90% were considered ‘transitional’ and those below 10% abundance were considered *Durusdinium* ‘negligible’.

### Whole genome sequencing and genotyping

Detailed information regarding DNA extraction methods, whole-genome sequencing and genotype calling can be found in ^37^. Briefly, libraries were prepared (Paired-end, 2 X 150bp, Kapa Hyper Prep) and sequenced (Illumina Novaseq X plus) by Admera Health Biopharma. Sequencing yielded an average 6.7 million paired end reads per sample (range: 4.5 - 14.8 million). All bioinformatic work was performed on the University of Southern California’s Center for Advanced Research Computing (CARC) system. Reads were trimmed to remove sequencing adapters, low quality ends (<Q20), and 5’ bias and filtered to retain those with a 99% base call accuracy over at least 90% of the read and mapped to a combined *A. palmata* (NCBI assembly: GCA_964030605.1) and Symbiodiniaceae genome reference (*Symbiodinium microadriaticum*^100^, *Breviolum minutum*^101^, *Cladocopium goreaui*^102^, *Durusdinium trenchii*^103^) and ambiguous reads/reads mapping exclusively to the symbiont references were discarded. Six samples were removed from the dataset due to low mapped read yields (< 2 million per sample which corresponded to low coverage (<1x) (Table S18). Mapping yielded an average coverage of 3.5x (range: 1-5x) for the 151 remaining samples. Variant call files (VCF) were generated per sample using the multiallelic caller in bcftools v1.21^104^ requiring a minimum map quality score of 40 and a minimum base quality score of 20 prior to merging. A population-specific haplotype panel and linkage map for *Acropora palmata*^105,106^ was used to recalibrate and impute genotype calls using a two-step imputation pipeline as detailed and validated in^37^. Note that one additional sample (AP66) was excluded from the sample set to be imputed due to its inclusion in the reference panel, leaving 150 shallow samples. In total, 1,615,451 variable sites were genotyped across samples, of which 131,065 were fully imputed.

### Genome Wide SNP Associations

A merged VCF of all unique experimental genets was created combining the 150 samples which underwent recalibration and imputation with independently generated high coverage whole genome sequence data for genet AP66 yielding a final whole genome genotyped set of 151 samples. However, trait data were missing for one genet from the 2020 biparental cross between two Lower Keys founder genets, AP358, and thus a total of 150 genets were used for the GWAS analysis. We focused on four traits (1) the proportion of *Durusdinium* hosted by a genet, (2) mean genet survival under heat stress, (3) mean genet surface area growth in the control treatment and (4) the proportional change in mean genet surface area growth in heat relative to the control treatment (Fig. 1D).

*Durusdinium* proportion was predominantly driven by genet identity (Fig. 2,S1) and the distribution was almost binary, therefore a quantile normalization of the continuous trait measure did not accurately reflect genet-specific differences in trait values (Fig. S15). We therefore coded *Durusdinium* dominance as a binary with 1 reflecting coral with >90% *Durusdinium* relative abundance and 0 reflecting coral genets with <90% abundance. Only certain outcomes were possible for mean genet mortality (e.g. 0, ⅓, ⅔, or all ramets died over the course of the experiment). We therefore also coded survival in the heat treatment as a binary with 1 reflecting coral with 100% ramet survival and 0 reflecting genets with any ramet mortality over the course of the experiment, which was equivalent to binning of genet-specific hazard ratios in heat-treated ramets. The final two traits, surface area growth in control and the proportional change in surface area growth in heat relative to control, were continuous distributions which we quantile normalized.

We tested associations between individual SNPs and each individual trait using linear mixed models as implemented in the software GEMMA v0.98.5^107^. For all models we controlled for genet relatedness using KING kinship coefficients^37^, as a comparison of relatedness estimates derived from GEMMA relative to the KING kinship coefficients revealed an asymptote (Fig. S16). Specifically, GEMMA relatedness estimates were negative for many individuals with confirmed second degree relationships^37^. Therefore, we used the KING kinship coefficients, with the addition of 151 self-comparisons set at the maximum observed kinship level (0.443) to fulfill the format requirements of GEMMA. The models for survival and surface area growth also included *Durusdinium* proportion (coded as a binary) as a covariate to account for the influence of symbiont type on holobiont trait values. For the input genotypes, we used high quality genotype calls in BED format. We ignored sites containing more than 10% missing individuals. Additionally, we filtered out SNPs with Hardy-Weinberg p values below 10^-7^, and with a minor allele frequency (MAF) less than 0.05 and randomly selected one representative SNP within tight linkage blocks (R^2^>0.97), leaving a total of 427,020 SNP correlations which were tested for their association with phenotype. We set the significance threshold as 5 x 10^-8^, a commonly used cutoff in human genetic studies, which was also found to be an appropriate threshold for coral^5^.

### Fine-mapping top candidate regions

To identify candidate genes potentially associated with peaks, we focused on SNPs that passed significance thresholds following GWAS. For *Durusdinium* retention and surface area growth in the control treatment, SNPs meeting the genome-wide significance threshold of P < 5 × 10□□ were retained. For proportional change in surface area growth under heat stress, we applied a genome-wide suggestive threshold of P < 1 × 10□□. SNPs located within 100 kb of one another were clustered and assigned to the same locus. This resulted in the identification of 7, 2, and 7 independent loci for the three traits respectively. To associate genomic regions with functional elements, we extracted all annotated genes located within ±100 kb flanking regions of each locus using a custom script. Gene positions and significant SNPs were visualized using the gggenes package in R. Furthermore, we conducted gene ontology (GO) enrichment analyses separately for each trait. For each set of candidate genes, we applied Fisher’s exact test using the GO_MWU package^108^, comparing the list of genes within associated loci against the full set of genes annotated in the genome.

To refine association signals identified in the GWAS, we conducted statistical fine-mapping with the susieR package (Wang et al. 2020). Fine-mapping analyses were restricted to genomic regions (±1 Mb) surrounding SNPs that surpassed the genome-wide significance threshold (P < 5 × 10^-8^). Fine mapping was conducted on a total of 1,143,642 SNPs without additional linkage disequilibrium (LD) pruning. We computed the LD matrix using PLINK v1.9 with the --r square option. GWAS summary statistics for these SNPs, including effect sizes and standard errors, were subset to match the LD matrix. SuSiE was then run using the susie_rss function. We used default settings with 10 effect components. Credible sets were constructed at a 95% confidence level. Locus plots showing association signals and PIP plots were generated in R using ggplot2.

### Melanocortin receptor homology and phylogenetic analysis

To evaluate whether genes annotated as melanocortin receptors in the *A. palmata* genome represent genuine homologs of the vertebrate melanocortin system, we assembled a dataset of candidate melanocortin receptor sequences spanning anthozoans and vertebrates. Specifically, reference MC4R and MC5R proteins were retrieved from NCBI for nine vertebrate model species spanning teleosts, amphibians, birds and mammals (Table S19) in addition to Human MC4R (NP_005903.2) and MC5R (NP_005904.1) protein sequences. These were used as blastp queries^109^ against the gene models of *Acropora tenuis*, *Pocillopora damicornis, Stylophora pistillata*, and the *Exaiptasia diaphana* CC7 genome from the Reef Genomics database^110^ and all sequences annotated as MC4R or MC5R in the *A. palmata* genome. Hits were retained at an e-value threshold of 1 × 10^−20^, and identical sequences were removed based on gene ids. In addition, human MC4R (NP_005903.2) and MC5R (NP_005904.1) protein sequences were used as blastp queries against all Symbiodiniaceae gene models from Reef Genomics database but no hits were retrieved, even with a relaxed e-value threshold of 1 × 10^−5^.

Amino acid sequences were aligned with MAFFT v7.520^111^ then trimmed with trimAl v1.4.1 under the automated1 heuristic^112^. A maximum likelihood phylogenetic tree was reconstructed with IQ-TREE 2^113^, with the best-fitting substitution model selected by ModelFinder. Branch support was assessed using 1,000 ultrafast bootstrap replicates.

*Gene expression analysis of* A. palmata *genets following symbiont community shuffling and surface area growth in response to a field transplantation*

To further investigate the functional role of candidate loci associated with *Durusdinium* retention and surface area growth, we leveraged an independent transplant experiment in which positive growth and variation among genets in initial endosymbiont associations and subsequent shuffling from *Durusdinium* to Symbiodinium dominated communities was also observed^39^. Full methods and ITS2 amplicon sequencing results are described in Elder et al. (2023). Briefly, three ramets of five exclusively ex situ nursery-reared *A. palmata* genets were transplanted to each of nine reefs in the lower Florida Keys (n = 27 ramets per genet) in 2018. Outplanted corals were monitored every three months for the first year and biannually thereafter for growth (using 3D photogrammetry^114^) and survival for a total of two years. *In situ* photographs of individual coral over time were used to generate 3D models of each ramet in Metashape 1.5.4 (Agisoft LLC, St. Petersburg, Russia) using a high-throughput pipeline^114^. Specifications for model building and all scripts can be found at https://github.com/wyattmillion/Coral3DPhotogram. Each 3D model was then imported into Meshlab v2020.6^115^ to measure surface area (SA) following^114^. The effects of genotype and site on the change in SA was evaluated for outplants at sites with sufficient survival (Fig. 1E; Bahia Honda was excluded due to complete mortality) using linear mixed effects models implemented with the lmer package^116^. SA was square-root transformed to better satisfy assumptions of linearity and homoscedasticity and modeled as a function of time-point, as well as genotype, site and the genotype by site interaction including scalar random effects of ramet to account for repeated measures, and the site-specific array to which the ramet was outplanted. Although genets differed in their initial SA (Fig. S17), there was no relationship between a ramet’s initial SA and its SA at subsequent time-points (Fig. S18), therefore this covariate was dropped from models to minimize overfitting. Significance of fixed effects for all models were evaluated using a type III analysis of variance (Kenward-Roger’s method) followed by Tukey’s pairwise comparisons to determine differences among factor levels when warranted.

Tissue samples were taken from a subset of 10-12 ramets per genet prior to outplanting (T0) and in surviving coral (N=92 ramets) following two years in the field (T24), flash frozen in liquid nitrogen, and stored at -80 C until processing following best practice guidelines for gene expression and microbiome sampling^117^. No *Symbiodinium fitti* were detectable prior to outplanting (T0) in most genotypes. Four genets were exclusively dominated by a *Durusdinium* (∼*trenchii*) ITS2 type, whereas genet 13-XK showed consistent co-infection with a *Cladocopium* ITS2 type composed of three co-occurring C1 amplicon sequence variants at T0 (Fig. 1E). Four sequence variants (0.1% of reads) matching to an *S. fitti* type were identified in one ramet of genet 13-XK at T0^39^. No bleaching was observed during quarterly site visits, but holobiont survival varied across sites^39^. *Durusdinium* remained the dominant ITS2 profile after 2 years of outplanting^39^ (T24, Fig. 1E). Surviving co-infected 13-XK ramets shuffled to *Durusdinium* dominance across sites and strains of the homologous symbiont, *Symbiodinium fitti*, were also detected in ramets of all genets at four sites^39^. Here, we focused on a subset of three sites (Dave’s Ledge, Eastern Dry Rocks, Looe Key, Fig. 1A,E) at which complete shuffling from Durusdinium (∼*trenchii*) to *Symbiodinium fitti* dominated endosymbiont communities was observed in some ramets^39^.

To profile global gene expression, tissue was removed from the skeleton and homogenized in Aurum Total RNA lysis buffer using the OMNI International bead rupture elite with sterile glass beads. Nucleic acids were extracted from all samples using the BioRad Aurum Total RNA Mini Kit following the manufacturer’s protocols, omitting the on-column DNAse step. The gene expression aliquot was DNAse treated using Invitrogen’s DNA-free DNA Removal Kit and 1 microgram of high quality RNA was used to prepare 3’ tag-based RNA-seq libraries ^118^ following standard protocols (https://github.com/z0on/tag-based_RNAseq). Libraries made with Kapa hyper prep minimal PCR were pooled in equimolar concentrations and sequenced on the Illumina NextSeq 500 platform (1x75bp) at the USC Genome Core. Sequencing yielded a total of 475.23 million raw reads across 142 samples. Per-sample library sizes ranged from 0.28 to 10.21 million reads, with a median of 3.06 million reads (Table S20). PCR duplicates and primer sequences specific to the Tag-Seq protocol were removed using custom perl scripts. Poly-A tails were clipped using fastx_clipper from FASTX-Toolkit v0.0.14^119^. Adapter contamination was removed using BBDuk from BBMap v39.26^120^. Quality filtering was then performed using fastq_quality_filter from FASTX-Toolkit v0.0.14 with parameters -q 20 -p 70 -Q33 which yielded a total of 303.00 million clean reads (0.19 to 6.53 million reads per sample, median = 1.98 million reads, Table S20). Quality-filtered reads were mapped to a concatenated host–symbiont reference genome using STAR v2.7.11b^121^ to enable assignment of reads to either host or symbiont origin. The reference genome consisted of the *A. palmata* assembly jaAcrPala1.3 (NCBI accession GCA_964030605.3) concatenated to the *Durusdinium trenchii* genome assembly SCF082 (NCBI accession GCA_963969995.1). Non-model species often have incomplete 3’ UTR annotations^122^, and 3’ tag-based RNA-seq (Tag-Seq) libraries are enriched for 3’ UTR transcripts^118^. Therefore gene models of *A. palmata* were extended to improve 3’ untranslated region (UTR) annotations following Levy et al. (2021). The percentage of uniquely mapped reads ranged from 31.68% to 83.17%, with a median of 69.89% across samples (Table S20). The resulting BAM files were used to define genomic intervals enriched for Tag-seq signal using MACS2^123^. Stranded genomic intervals were used to extend the annotated 3’ ends of gene models. Gene-level counts were quantified using featureCounts v2.1.1^124^ with the extended gene annotation. Of 24,099 total gene models in the *A. palmata* jaAcrPala1.3 genome assembly, 16,592 genes (68.85%) were extended and the mean extension length was 1,022 bp. Gene-level quantification using featureCounts yielded assignment rates ranging from 33.16% to 70.36%, with a median of 64.25% across samples. The mean improvement in assignment rate compared to using the original gene models was 28.97% (from 34.45% to 63.42%, Table S20). Genes with total counts fewer than 10 across all samples were excluded prior to downstream statistical analysis. Principal component analysis was performed on normalized count data. PCA plots were generated using ggplot2 v3.5.2^125^ to assess global expression patterns. Differential expression analysis was performed using DESeq2 1.46.0^126^ using a series of contrasts described below.

Prior to transplantation, four genotypes were exclusively dominated by *Durusdinium*, whereas genotype 13-XK exhibited co-infection with *Cladocopium*^39^. After two years of outplanting, some ramets remained dominated by *Durusdinium*, while others became dominated by *Symbiodinium*^39^. For tissue samples collected from the three focal reef sites at T24, ramets were classified according to symbiont composition using two categorical variables: presence of *Symbiodinium* (Y/N) and dominant symbiont type (A = *Symbiodinium* dominated; D = *Durusdinium* dominated). Associations between genotype and symbiont category were tested using Fisher’s Exact Test implemented in R with the function fisher.test. Differential expression analysis was then performed including ramets of all five genets. Samples were grouped according to dominant symbiont type (A vs. D) and by presence versus absence of Symbiodinium (Y vs. N). Genes were considered differentially expressed if they met both the criteria |log2 FC| > 1 and adjusted p-value < 0.05. This method identified gene expression differences associated with Symbiodinium dominance (A vs. D) and Symbiodinium presence (Y vs. N) following shuffling (Fig. 1F).

We also modeled expression as a function of SA growth. Instances of negative growth attributable to predation by *Drupella* snails were removed before expression-growth modeling. Specifically, matched gene-expression samples were excluded if T0 or T24 surface-area measurements were missing, or if any observed adjacent interval from T12 to T24 showed a surface-area decline greater than 10%. Missing intermediate timepoints were ignored for breakage calling, but T0 and T24 measurements were required for inclusion. This filtering excluded seven matched gene-expression samples with predation losses resulting in retention of 80 samples for the final expression-growth analysis. Relative SA growth was calculated for remaining ramets as (T24 SA − T0 SA) / T0 SA. Low expression transcripts were excluded from modeling if expression was zero in at least 90% of retained samples; after this filter, 18,222 of 28,920 transcripts were tested. For each retained transcript, linear models were fit with expression as the response variable and relative surface-area growth, field site, and dominant symbiont category as predictors, including 1) expression ∼ rel_growth, 2) expression ∼ location, 3) expression ∼ location + rel_growth, and 4) expression ∼ location + dom_symb + rel_growth. Term-level F tests were extracted from each model, and p-values were adjusted using the Benjamini-Hochberg false discovery rate (FDR) procedure separately within each model-term combination. Transcripts were first considered growth-associated if the relative-growth term in the expression ∼ relative growth model had FDR < 0.05. This initial screen identified 54 transcripts. To determine whether these associations remained after accounting for field site, each growth-associated transcript was then tested with a nested ANOVA comparing expression ∼ field site to expression ∼ field site + relative growth. Follow-up ANOVA p-values were Benjamini-Hochberg adjusted across the 54 growth-associated transcripts, and transcripts with follow-up FDR < 0.05 were retained as the final location-controlled surface-area growth gene set. This final analysis identified 49 transcripts associated with relative surface-area growth. Gene Ontology (GO) enrichment analysis was conducted using the GO_MWU package but no terms remained after multiple test correction.

Finally, we integrated results from the GWAS and DEG analyses to identify genes supported by both association and expression analyses. Gene lists in association with previously identified GWAS loci were refined to include only genes in proximity to significant credible sets (Table S10). Intersections were then performed between these credible set cc_genes and the two sets of DEGs respectively: those differentially regulated based on either the presence or dominance of *Symbiodinium fitti* following field transplantation (Table S14) and those correlated with surface area growth (Table S15).

## ACKNOWLEDGEMENTS

We thank Eleftherios Karabelas for contributing to the execution of the project and primary data collection, Daniella Leon for help with DNA extractions and Erich Bartels for fieldwork support. Work was conducted under permits FKNMS-2021-172 and SAL-22-2406-SCRP to Mote Marine Laboratory. Funding for the GWAS experiment was provided by NOAA Ruth D. Gates Coral Restoration Innovation Grant #NA21NMF4820300 to CDK and EMM. The field transplant experiment was supported by private funding from the Alfred P. Sloan and Rose Hills Foundation to CDK and conducted under permits FKNMS-2015-163-A1 and FKNMS-2018-035. Support for additional functional genomic analyses was provided by the Allen Family Philanthropies to CDK.

## DATA ACCESSIBILITY

Raw sequence data have been archived on NCBI’s SRA under PRJNA1078365. Scripts necessary to replicate bioinformatic processing steps, subsequent statistical analyses and data visualizations for analysis of trait data and the GWAS can be found at https://github.com/ckenkel/ApalGWAS while those for fine mapping and the gene expression analysis can be found at https://github.com/Ruiqi-CUB/AcrPal_GWAS_GitHub (Zenodo DOIs to be generated).

