## Supplemental Tables and Figures for "Genomic variation associated with endosymbiont shuffling and areal growth in the endangered elkhorn coral, *Acropora palmata*"

[1]University of Southern California, 3616 Trousdale Parkway, Los Angeles, CA 90089, USA
[2]Australian Institute of Marine Science, PMB3, Townsville MC QLD 4810, Australia
[3]Mote Marine Laboratory, Sarasota, FL 34236, USA
[4]University of California, Santa Barbara, CA 93106, USA
[5]Department of Biology, Hopkins Marine Station, Stanford University, Pacific Grove, CA 93940
[6]Department of Integrative Biology, Oregon State University, Corvallis, OR 97331
[7]Department of Marine Biology and Ecology, Rosenstiel School of Marine, Atmospheric, and Earth Science, University of Miami, Miami, Florida 33149
[8]University of Colorado Boulder
[9]DLNR - Division of Aquatic Resources, 74-380B Kealakehe Pkwy, Kailua Kona, HI 96740
[10]Florida Fish and Wildlife Conservation Commission, 2796 Overseas Highway, Marathon, Florida, 33050

**Figure S1**. Proportional abundance of *Durusdinium (~trenchii)*, *Symbiodinium (~fitti)* and *Cladocopium* spp. within focal *Acropora palmata* genets by family origin and temperature treatment. Genet order along the x-axis is identical across panels and reflects rank order according to the mean proportion of *Durusdinium* in control samples. Note that while naming conventions changed at IC2R3 over time, AP20 (E-H-SI Batch Cross) and 13-XK (Fig 1F) are ramets of the same genet and *Cladocopium* spp. was detected in the IC2R3 nursery both here and by Elder et al. (2023).

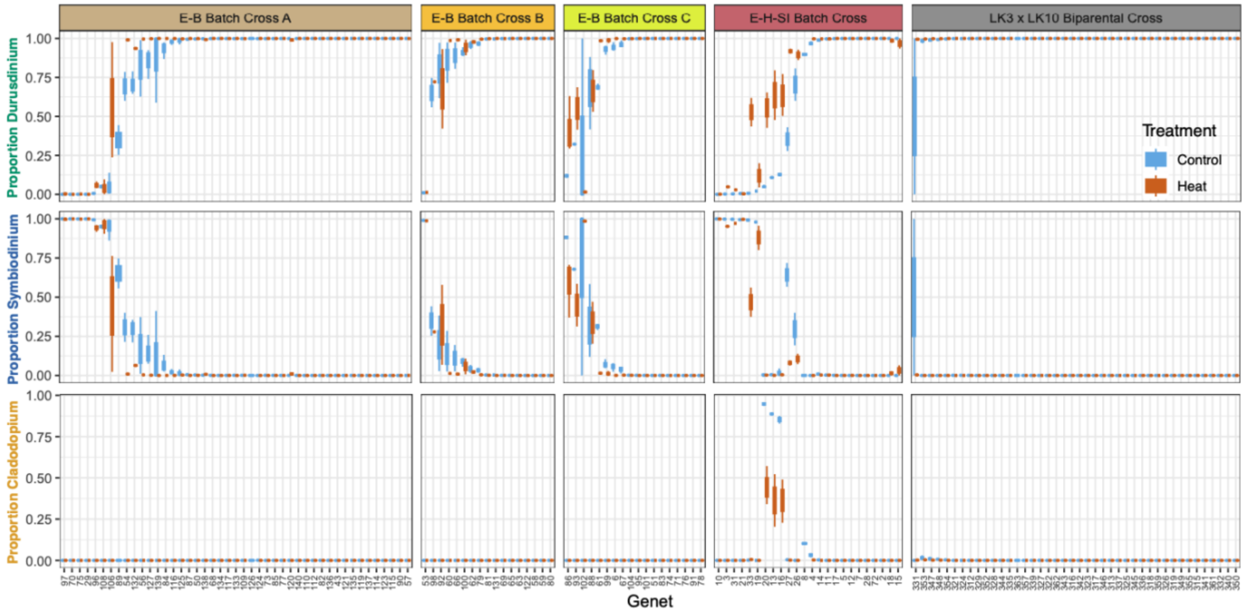


**Table S1**. Summary of linear mixed effects model assessing the effect of the proportion of *Durusdinium*, as well as family origin, treatment and their interaction on log10 transformed S:H cell ratio as implemented in the lme4 package.

Type III Analysis of Variance Table with Kenward-Roger's method

| | Sum Sq | Mean Sq | NumDF | DenDF | F value | Pr(>F) | |
|---|---|---|---|---|---|---|---|
| X.D | 5.0890 | 5.0890 | 1 | 165.00 | 17.0233 | 5.844e-05 | *** |
| Origin_PostHoc | 4.9766 | 1.2442 | 4 | 140.32 | 4.1617 | 0.003239 | ** |
| Treatment | 0.5672 | 0.5672 | 1 | 4.18 | 1.8973 | 0.237492 | |
| Origin_PostHoc:Treatment | 0.6307 | 0.1577 | 4 | 571.04 | 0.5274 | 0.715625 | |

---

Signif. codes: 0 '***' 0.001 '**' 0.01 '*' 0.05 '.' 0.1 ' ' 1

**Figure S2**. Distribution of genet S:H cell ratios as a function of family origin and the mean proportion of *Durusdinium* hosted by a genet in heat treated ramets. Genet order is identical to that in Figure 2/S1 and reflects the mean proportion of *Durusdinium* in control ramets.

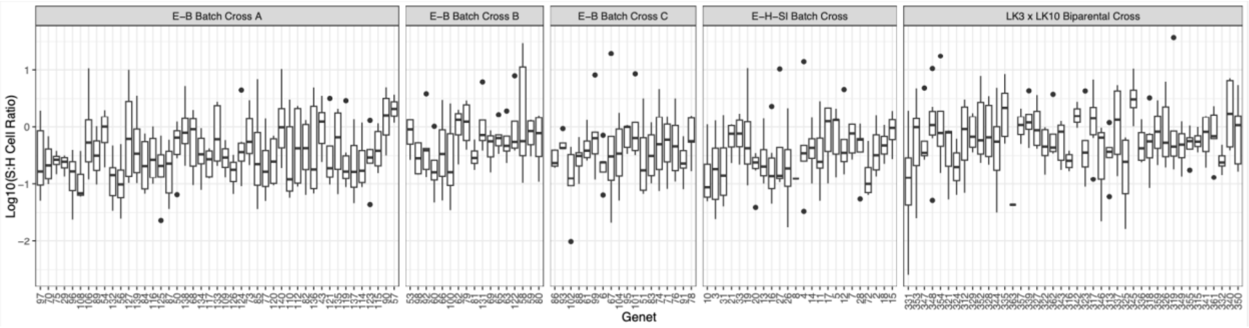


**Table S2**. Summary of linear mixed effects model assessing the effect of the proportion of *Durusdinium*, as well as family origin, treatment and their interaction on the proportional change in Fv/Fm as implemented in the lme4 package.

```
Type III Analysis of Variance Table with Kenward-Roger's method
                         Sum Sq Mean Sq NumDF  DenDF F value    Pr(>F)
X.D                     0.65426 0.65426     1 213.19  49.097 3.147e-11 ***
Origin_PostHoc          0.63381 0.15845     4 145.86  11.889 2.183e-08 ***
Treatment               0.09223 0.09223     1   4.08   6.921 0.0569294 .
Origin_PostHoc:Treatment 0.27766 0.06941    4 500.06   5.209 0.0004062 ***
---
Signif. codes:  0 ‘***’ 0.001 ‘**’ 0.01 ‘*’ 0.05 ‘.’ 0.1 ‘ ’ 1
```

**Figure S3**. Distribution of proportional change in Fv/Fm values as a function of family origin and the mean proportion of *Durusdinium* hosted by a genet. Genet order along the x-axis is identical to that in Figure 2/S1 and is ordered left to right based on the mean proportion of *Durusdinium* in control ramets (~low to high).

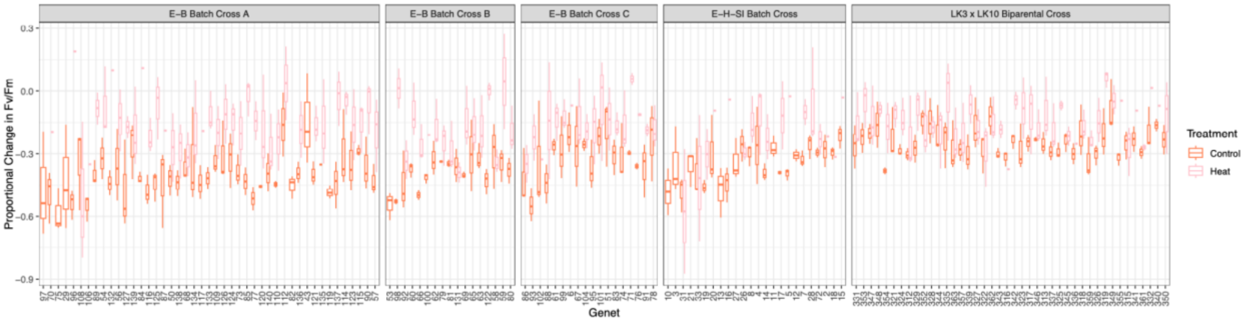

**Figure S4**. Partial mortality by family origin and treatment. Points are colored by instances of no partial mortality (grey) vs a positive change in the planar surface area of dead skeleton (red) by cross type and treatment (~zero-inflated model).

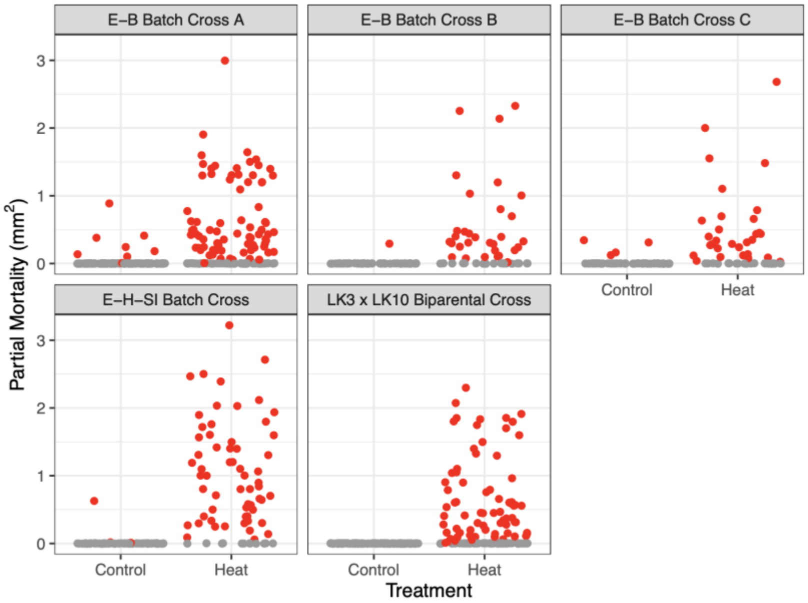


**Table S3**. Summary of zero-inflation generalized linear mixed model assessing the effect of the proportion *Durusdinium*, family origin, and treatment on partial mortality as implemented in the glmmTMB package.

**Random effect variances**

| | Groups | Name | Std.Dev. |
|---|---|---|---|
| 1 | Raceway | (Intercept) | 2.021e-05 |

**Conditional fixed effects**

| | Estimate | Std. Error | z value | Pr(>\|z\|) |
|---|---|---|---|---|
| (Intercept) | 4.10 | 1.13 | 3.65 | 0.00 |
| X.D | 0.76 | 0.21 | 3.55 | 0.00 |
| Origin_PostHocE-B Batch Cross A | -0.78 | 0.50 | -1.56 | 0.12 |
| Origin_PostHocE-B Batch Cross C | -0.54 | 0.58 | -0.93 | 0.35 |
| Origin_PostHocE-H-SI Batch Cross | -1.29 | 0.49 | -2.63 | 0.01 |
| Origin_PostHocLK3 x LK10 Biparental Cross | -1.09 | 0.50 | -2.17 | 0.03 |
| TreatmentHeat | -2.34 | 1.01 | -2.31 | 0.02 |

**Conditional zero-inflation effects**

| | Estimate | Std. Error | z value | Pr(>\|z\|) |
|---|---|---|---|---|
| (Intercept) | 3.20 | 0.56 | 5.71 | 0.00 |
| X.D | 0.39 | 0.41 | 0.93 | 0.35 |
| Origin_PostHocE-B Batch Cross A | 0.06 | 0.39 | 0.14 | 0.89 |
| Origin_PostHocE-B Batch Cross C | -0.03 | 0.47 | -0.07 | 0.94 |
| Origin_PostHocE-H-SI Batch Cross | -0.75 | 0.46 | -1.63 | 0.10 |
| Origin_PostHocLK3 x LK10 Biparental Cross | 0.37 | 0.40 | 0.93 | 0.35 |
| TreatmentHeat | -4.05 | 0.33 | -12.28 | 0.00 |

**Figure S5**. Partial mortality as a function of family origin and the mean genet proportion of *Durusdinium* in heat treated ramets. Boxplot distributions are restricted to cases where some partial mortality was observed (~conditional model) whereas points show all data colored by instances of no partial mortality (grey) vs an areal loss of tissue (red) by cross type. Genet order is identical to that in Figure 2/S1 and reflects the mean proportion of *Durusdinium* in control ramets.

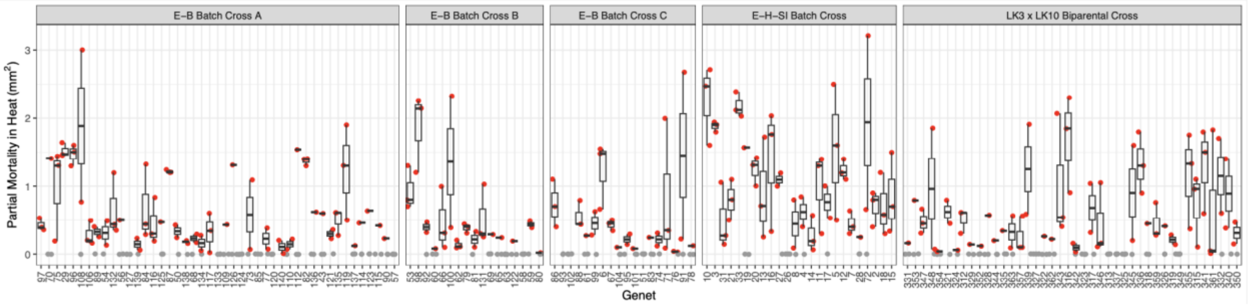


**Table S4**. Summary of zero-inflation generalized linear mixed model assessing the effect of the proportion *Durusdinium* and family origin on partial mortality in heat-treated ramets as implemented in the glmmTMB package.

**Random effect variances**

| | Groups | Name | Std.Dev. |
|---|---|---|---|
| 1 | Raceway | (Intercept) | 2.0903e-05 |

**Conditional fixed effects**

| | Estimate | Std. Error | z value | Pr(>\|z\|) |
|---|---|---|---|---|
| (Intercept) | 1.79 | 0.50 | 3.60 | 0.00 |
| X.D | 0.76 | 0.21 | 3.64 | 0.00 |
| Origin_PostHocE-B Batch Cross A | -0.78 | 0.50 | -1.58 | 0.11 |
| Origin_PostHocE-B Batch Cross C | -0.57 | 0.58 | -0.98 | 0.33 |
| Origin_PostHocE-H-SI Batch Cross | -1.32 | 0.49 | -2.71 | 0.01 |
| Origin_PostHocLK3 x LK10 Biparental Cross | -1.12 | 0.50 | -2.24 | 0.03 |

**Conditional zero-inflation effects**

| | Estimate | Std. Error | z value | Pr(>\|z\|) |
|---|---|---|---|---|
| (Intercept) | -1.35 | 0.62 | -2.17 | 0.03 |
| X.D | 0.89 | 0.53 | 1.68 | 0.09 |
| Origin_PostHocE-B Batch Cross A | 0.18 | 0.42 | 0.42 | 0.67 |
| Origin_PostHocE-B Batch Cross C | 0.31 | 0.50 | 0.62 | 0.53 |
| Origin_PostHocE-H-SI Batch Cross | -0.92 | 0.54 | -1.71 | 0.09 |
| Origin_PostHocLK3 x LK10 Biparental Cross | 0.26 | 0.43 | 0.62 | 0.54 |

**Table S5**. Summary of zero-inflation generalized linear mixed model assessing the effect of the proportion *Durusdinium*, as well as family origin, treatment and their interaction on the maximal change in planar surface area as implemented in the glmmTMB package.

**Random effect variances**

| | Groups | Name | Std.Dev. |
|---|---|---|---|
| 1 | Raceway | (Intercept) | 0.46597 |

**Conditional fixed effects**

| | Estimate | Std. Error | z value | Pr(>\|z\|) |
|---|---|---|---|---|
| (Intercept) | 2.36 | 0.52 | 4.56 | 0.00 |
| X.D | 0.20 | 0.39 | 0.51 | 0.61 |
| Origin_PostHocE-B Batch Cross A | 1.86 | 0.42 | 4.44 | 0.00 |
| Origin_PostHocE-B Batch Cross C | 2.00 | 0.56 | 3.56 | 0.00 |
| Origin_PostHocE-H-SI Batch Cross | 1.16 | 0.46 | 2.54 | 0.01 |
| Origin_PostHocLK3 x LK10 Biparental Cross | 0.44 | 0.36 | 1.23 | 0.22 |
| TreatmentHeat | 0.85 | 0.66 | 1.30 | 0.19 |
| Origin_PostHocE-B Batch Cross A:TreatmentHeat | -0.67 | 0.70 | -0.96 | 0.34 |
| Origin_PostHocE-B Batch Cross C:TreatmentHeat | -1.05 | 0.90 | -1.16 | 0.25 |
| Origin_PostHocE-H-SI Batch Cross:TreatmentHeat | 1.84 | 1.03 | 1.79 | 0.07 |
| Origin_PostHocLK3 x LK10 Biparental Cross:TreatmentHeat | 0.32 | 0.66 | 0.48 | 0.63 |

**Conditional zero-inflation effects**

| | Estimate | Std. Error | z value | Pr(>\|z\|) |
|---|---|---|---|---|
| (Intercept) | -4.61 | 0.88 | -5.23 | 0.00 |
| X.D | -0.02 | 0.47 | -0.04 | 0.97 |
| TreatmentHeat | 1.73 | 0.37 | 4.67 | 0.00 |
| Origin_PostHocE-B Batch Cross A | 1.15 | 0.77 | 1.50 | 0.13 |
| Origin_PostHocE-B Batch Cross C | 1.44 | 0.81 | 1.78 | 0.07 |
| Origin_PostHocE-H-SI Batch Cross | 1.84 | 0.78 | 2.34 | 0.02 |
| Origin_PostHocLK3 x LK10 Biparental Cross | -0.58 | 0.93 | -0.63 | 0.53 |

**Table S6**. Chi-square test of conditional model for surface area growth (i.e. only cases where some growth was observed)

```
Analysis of Deviance Table (Type II Wald chisquare tests)

Response: SA_dif_cor
                          Chisq Df Pr(>Chisq)
X.D                      0.2567  1    0.61238
Origin_PostHoc          30.6996  4  3.525e-06 ***
Treatment                3.3905  1    0.06557 .
Origin_PostHoc:Treatment 9.0449  4    0.05999 .
---
Signif. codes:  0 '***' 0.001 '**' 0.01 '*' 0.05 '.' 0.1 ' ' 1
```

**Figure S6**. Distribution of daily weight gain by family origin.

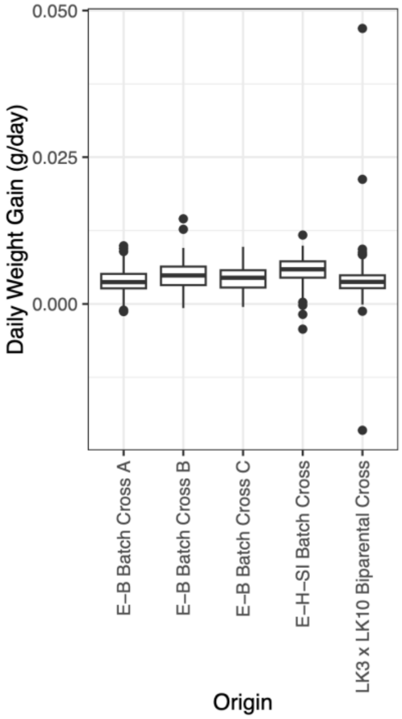


**Table S7**. Summary of linear mixed effects model assessing the effect of the proportion of *Durusdinium*, as well as cross origin, treatment and their interaction on daily weight gain as implemented in the lme4 package.

```
Type III Analysis of Variance Table with Kenward-Roger's method
                            Sum Sq    Mean Sq NumDF  DenDF F value    Pr(>F)    
X.D                     2.9170e-06 2.9170e-06     1 635.05  0.3584    0.5496    
Origin_PostHoc          3.0022e-04 7.5056e-05     4 634.75  9.2225 3.014e-07 ***
Treatment               2.3305e-05 2.3305e-05     1   5.08  2.8637    0.1505    
Origin_PostHoc:Treatment 7.8760e-06 1.9690e-06     4 634.77  0.2420    0.9145    
---
Signif. codes:  0 '***' 0.001 '**' 0.01 '*' 0.05 '.' 0.1 ' ' 1
```

**Figure S7**. Genome-wide association of propensity to retain *Durusdinium*. Boxplot distributions show individual genet trait values binned by genotype dosage and family origin. Horizontal lines on Manhattan plot indicate genome-wide suggestive threshold (blue, $P < 1 \times 10^{-5}$) and significant threshold (red, $P < 5 \times 10^{-8}$).

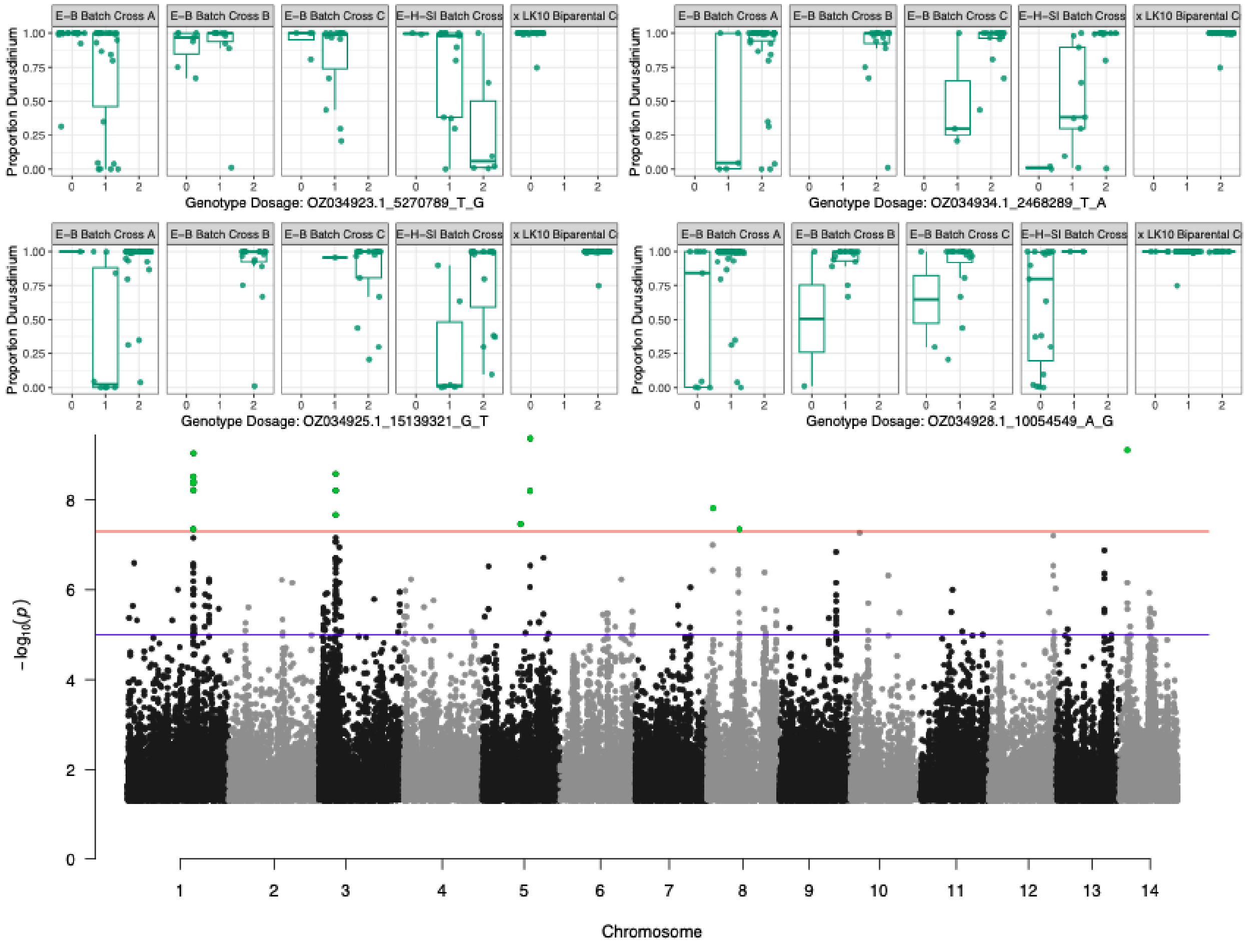

**Figure S8.** Maximum-likelihood phylogenetic tree of melanocortin receptor (MC4R and MC5R) homologs across cnidarians and vertebrates. The GWAS candidate gene g7717 from *A. palmata* MC4R is highlighted in blue (upper box); the reciprocally monophyletic vertebrate MC4R/MC5R clade is highlighted in lower blue box.

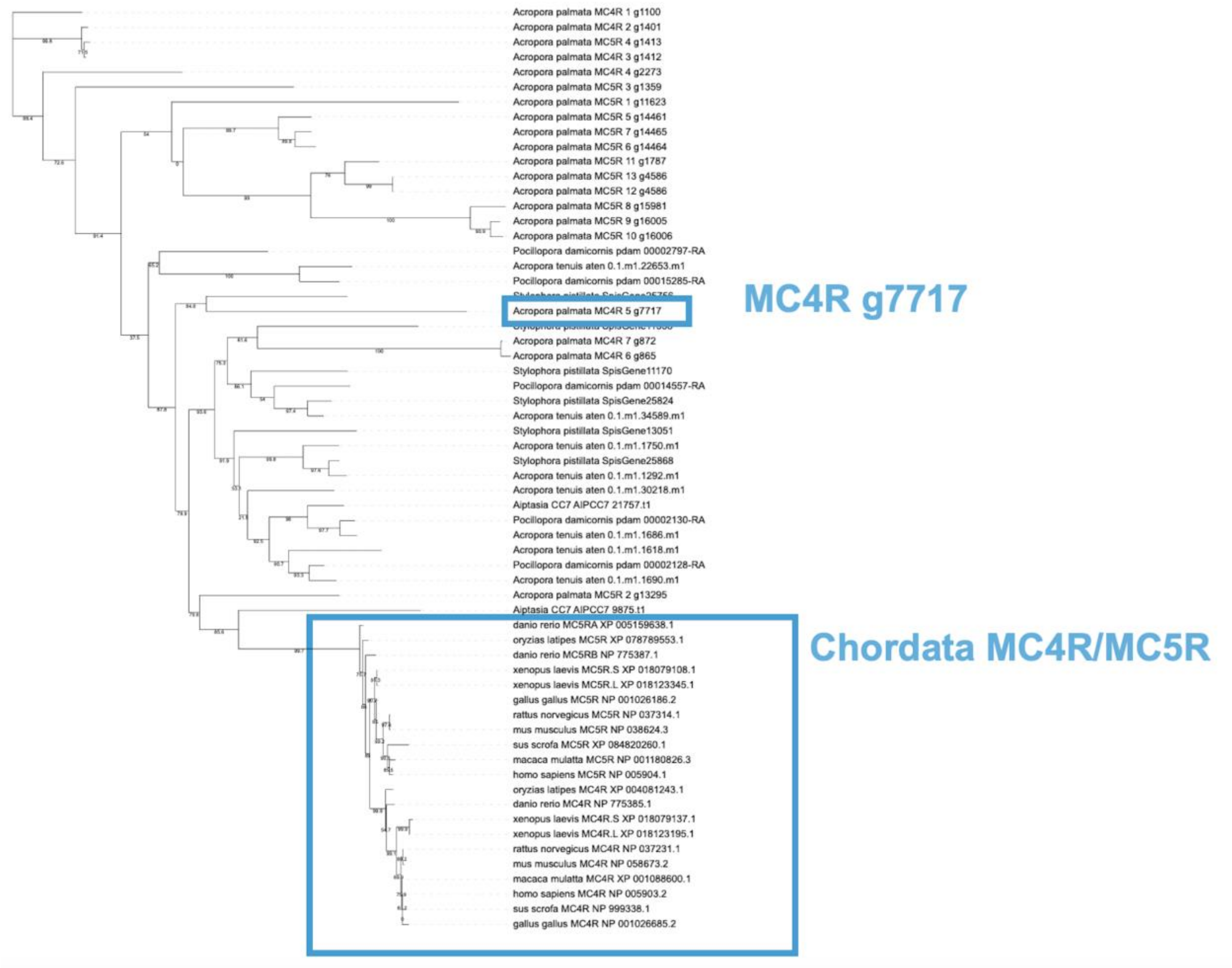


**Figure S9**. Genome-wide association of surface area growth in the control treatment. Boxplot distributions show individual genet trait values binned by genotype dosage and family origin.

Horizontal lines on Manhattan plot indicate genome-wide suggestive threshold (blue, $P < 1 \times 10^{-5}$) and significant threshold (red, $P < 5 \times 10^{-8}$).

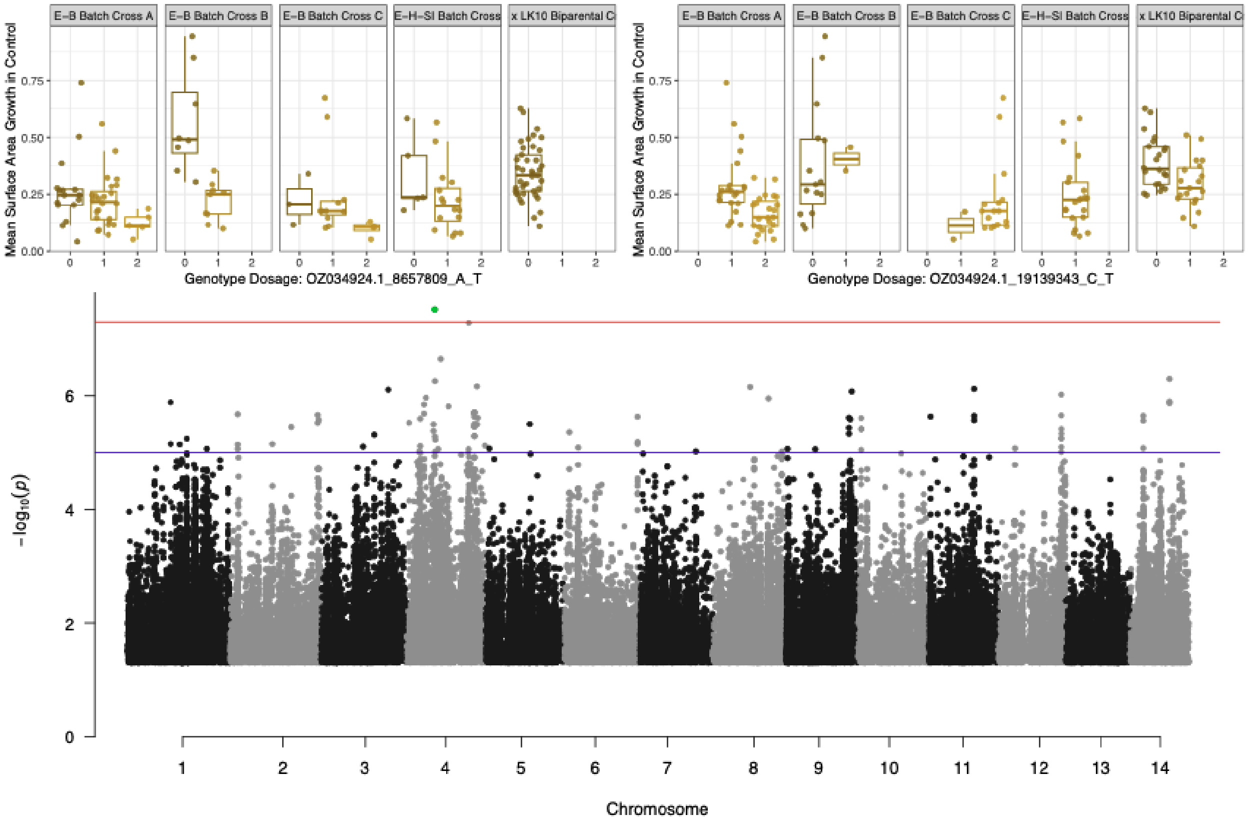


**Figure S10**. Genome-wide association of ramet mortality in heat treatment. Boxplot distributions show individual genet trait values binned by genotype dosage and family origin. Horizontal lines on Manhattan plot indicate genome-wide suggestive threshold (blue, $P < 1 \times 10^{-5}$) and significant threshold (red, $P < 5 \times 10^{-8}$).

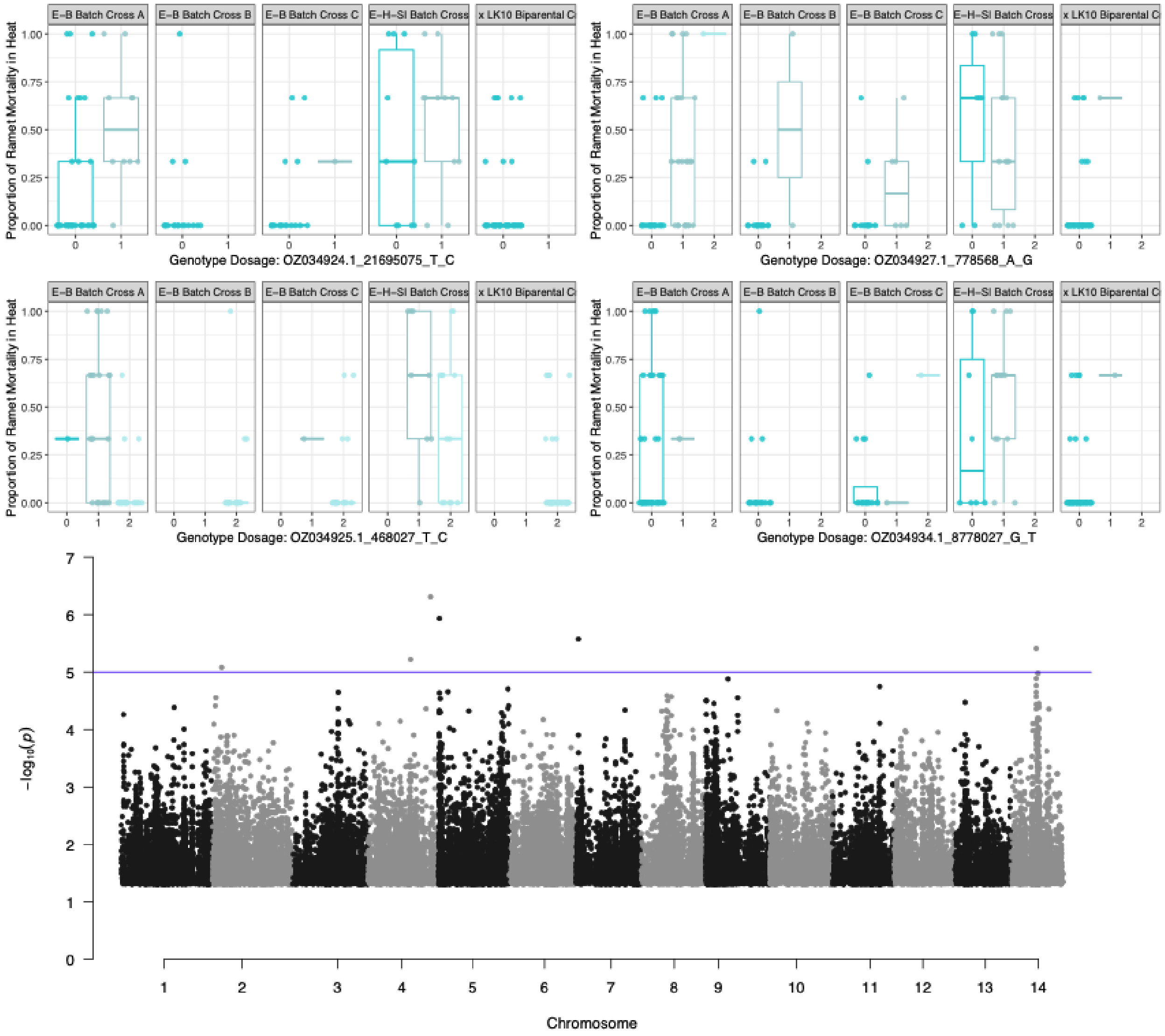


**Figure S11**. Genome-wide association of the proportional change in genet surface area growth in heat relative to the control treatment. Boxplot distributions show individual genet trait values binned by genotype dosage and family origin. Horizontal lines on Manhattan plot indicate genome-wide suggestive threshold (blue, $P < 1 \times 10^{-5}$).

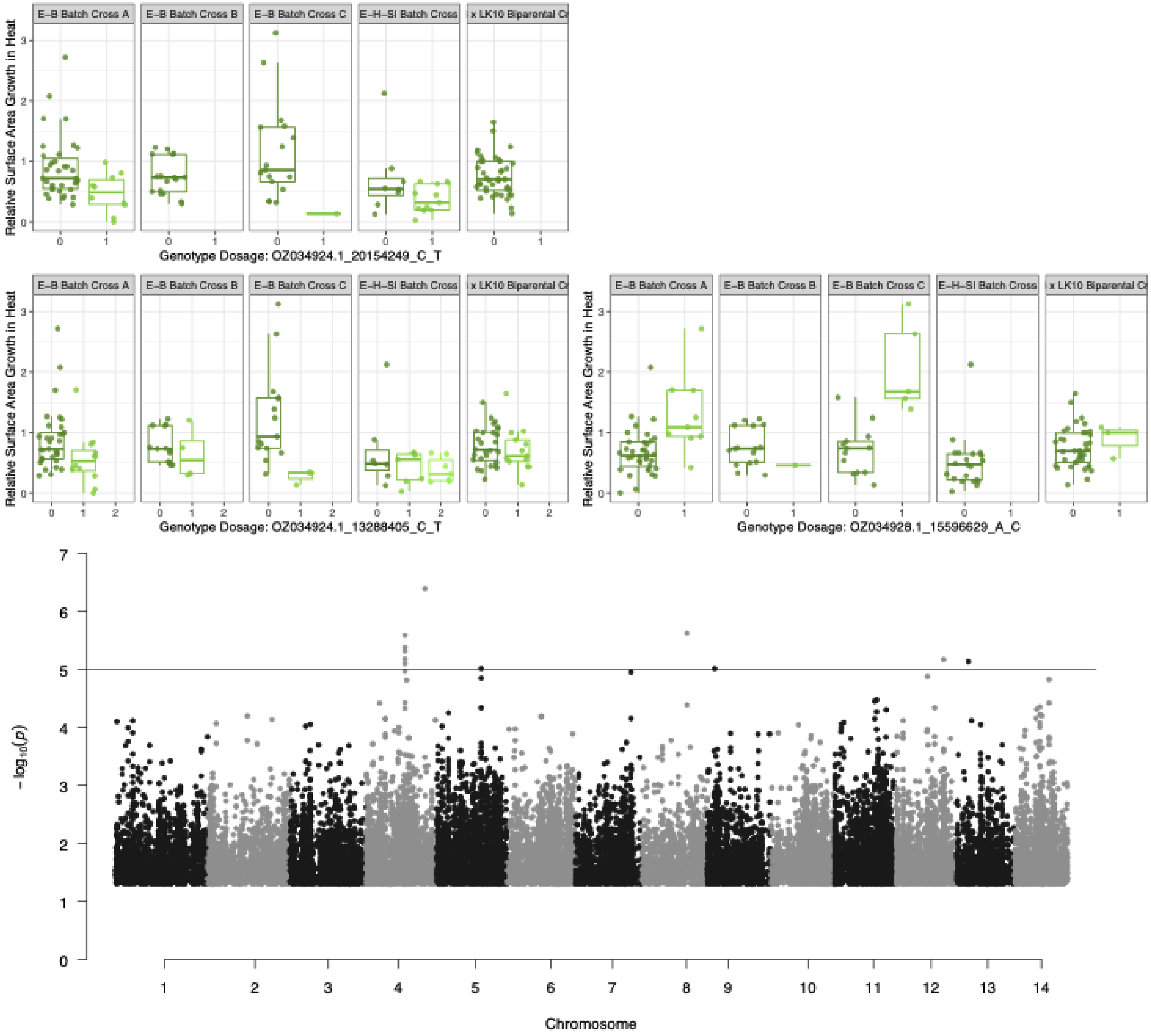


**Figure S12**. Mosaic plot of genotypes for three field transplant genets included in shallow whole-genome dataset (13-X5=G5, 13-X7=G7, 13-XK=G20, Fig. 1F,2) at chromosome 1 peak showing strongest association with propensity to retain *Durusdinium* (Fig. 3) by the proportion of ramets in which *Symbiodinium* (~*fitti*) was detected (present) or not (absent) at T24 (Elder et al. 2023).

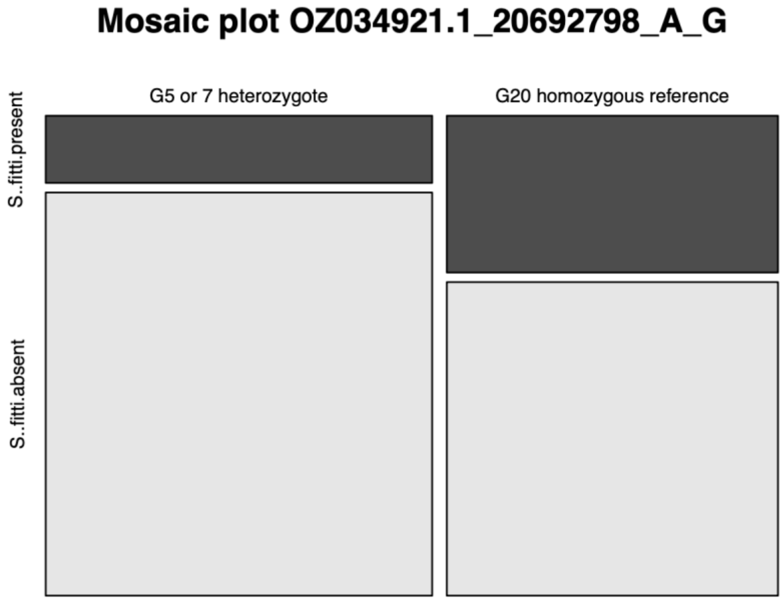


**Figure S13**. Boxplot distribution of proportion *Durusdinium* by genotype dosage and family origin for locus associated with surface area growth but in proximity to differentially expressed copy of MC4R in coral which shuffled to *Symbiodinium fitti* in the field.

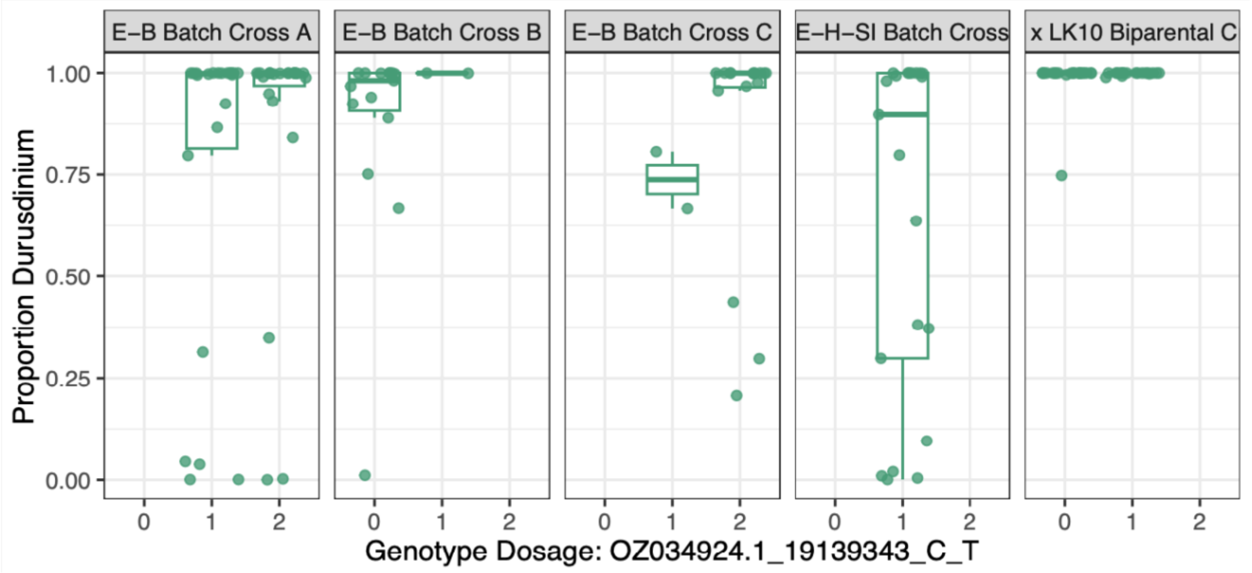


**Figure S14**. Amplification cycle (Cq) vs concentration for each symbiont to host cell dilution series

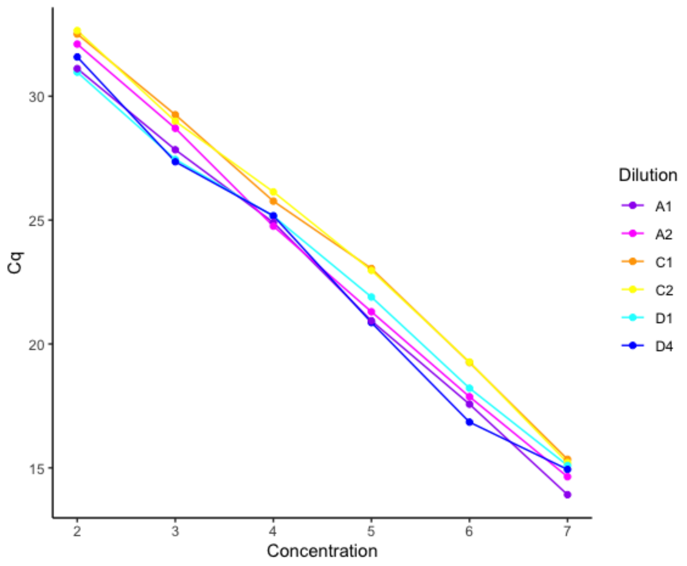


**Figure S15**. Histogram of mean and quantile normalized proportion of *Durusdinium* and correlation between raw and transformed trait values.

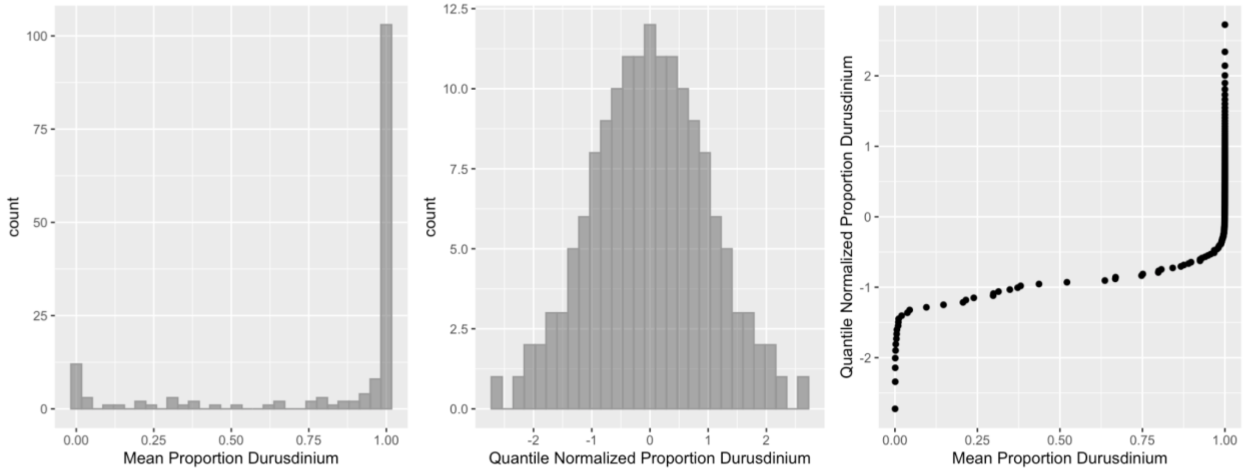


**Figure S16**. Correlation between GEMMA-based pairwise relatedness and KING kinship coefficients.

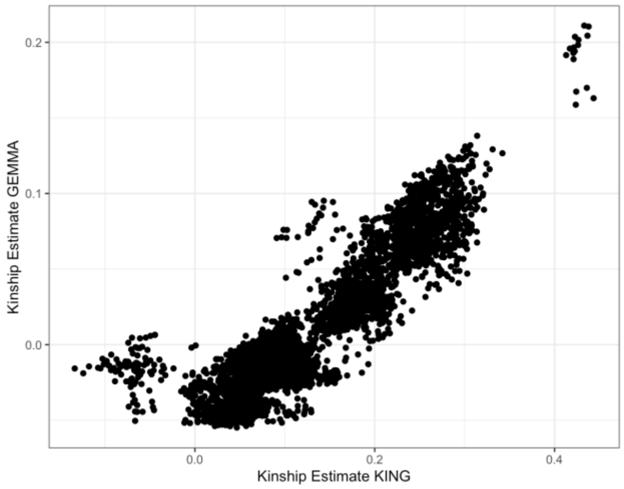


**Figure S17**. Boxplot distribution of initial surface area ($cm^2$) of outplanted ramets grouped by genotype.

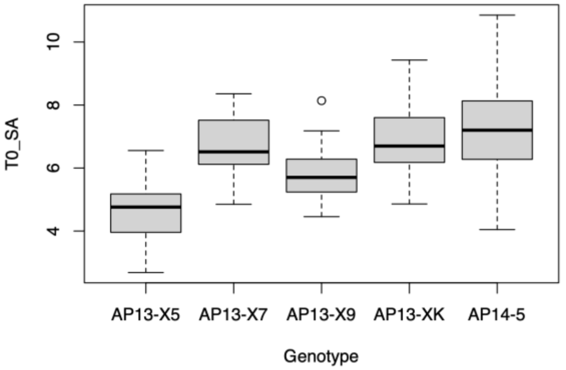


**Figure S18**. Ramet surface area at each monitoring timepoint post-outplanting as a function of initial surface area.

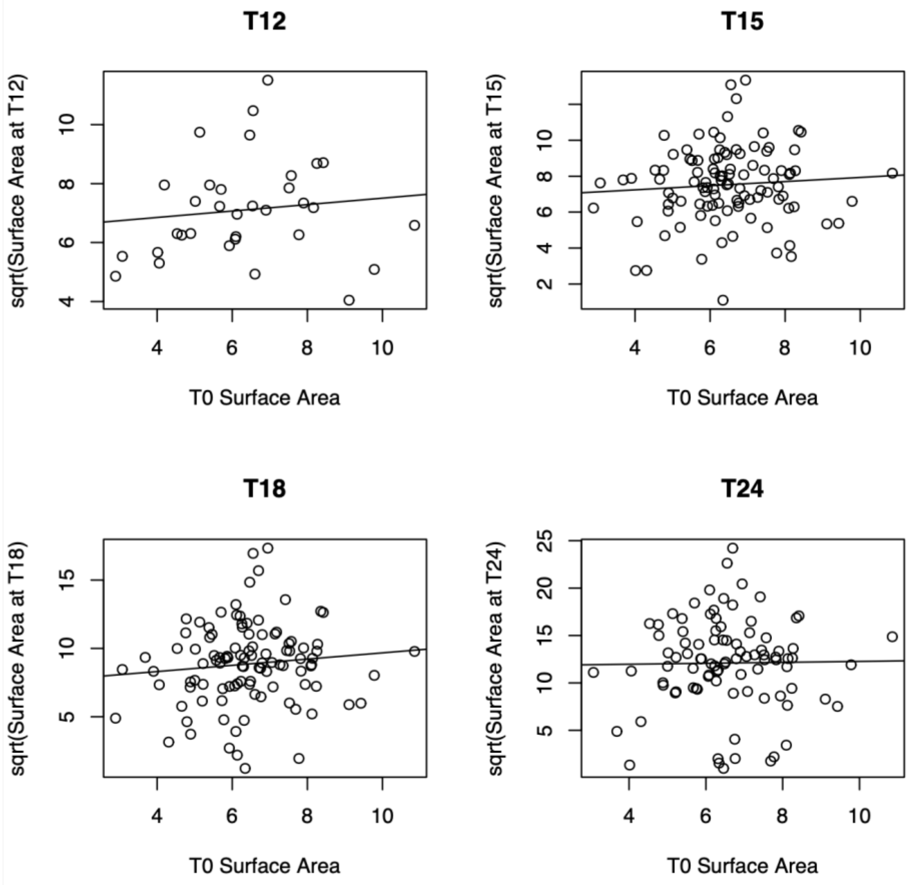
T12
T15
T18
T24
sqrt(Surface Area at T12)
sqrt(Surface Area at T15)
sqrt(Surface Area at T18)
sqrt(Surface Area at T24)
T0 Surface Area
T0 Surface Area
T0 Surface Area
T0 Surface Area
